# Structure of *Toxoplasma gondii* glideosome-associated connector suggests a role as an elastic element in actomyosin force generation for gliding motility

**DOI:** 10.1101/2022.12.09.519741

**Authors:** Yu-Fu Hung, Qu Chen, Isa Pires, Peter B. Rosenthal, Inari Kursula

## Abstract

*Toxoplasma gondii* glideosome-associated connector (GAC) is a giant armadillo-repeat protein, essential for parasite motility and conserved across Apicomplexa. It connects actin filaments to the plasma membrane *via* interactions with phosphatidic acid and membrane-spanning adhesins. It is unclear how GAC contributes to gliding motility and invasion and why such a large connector is needed. We determined the crystal structure of full-length *T. gondii* GAC at 2.3 Å resolution and explored its conformational space in solution using small-angle X-ray scattering and cryogenic electron microscopy. The crystal structure reveals a compact conformation but, in solution, GAC adopts both compact and extended forms. The PH domain stabilizes the compact form and may act as a switch triggered by membrane sensing. Based on its spring-like architecture, we suggest a role for GAC as an elastic element in actomyosin force generation during gliding motility and invasion.

---

The diseases malaria and toxoplasmosis are caused by obligatory intracellular parasites of the apicomplexan phylum, *Plasmodium* spp. and *Toxoplasma gondii* (*Tg*), respectively. These parasites display a unique substrate-dependent form of motility, termed gliding, during the motile and invasive stages of their life cycle^1^. Force for gliding motility is generated by the parasite actomyosin system^2^, and the whole gliding machinery is bridged from the inner membrane complex (IMC) to the parasite plasma membrane through a connection between actin filaments and adhesins formed by a 280 kDa protein called glideosome-associated connector (GAC)^3^. GAC is conserved throughout Apicomplexa, unique to these parasites, and essential for gliding motility^3^. The N-terminal part of GAC binds to and stabilizes actin filaments, the middle part is required for conoid targeting, and the C-terminal part associates with adhesins in the parasite plasma membrane^3^. A pleckstrin-homology (PH) domain in the C terminus binds to phosphatidic acid (PA) enriched membranes^3^.

Preliminary structural analysis and homology modeling of GAC suggested a club-shaped molecule with a maximum dimension of 27 nm for the full-length (FL) *Tg*GAC and 16 nm for the N-terminal actin-filament-binding region^3^. FL-*Tg*GAC was predicted to possess a substantial armadillo-repeat region with a wider end around the actin-binding region and a narrow end with the PH domain at the tip, far from the N-terminal actin-binding region^3^. Armadillo-repeat domains are in general involved in protein-protein interactions^4^. GAC is a unique example of an armadillo-super-helical structure directly binding to and stabilizing filamentous actin^3^. However, its exact actin-binding mode and function in the glideosome are unknown.

Here, we present a high-resolution crystal structure of *Tg*GAC and explore its conformational space using single-particle cryogenic electron microscopy (cryo-EM) and small-angle X-ray scattering (SAXS). The crystal structure reveals a surprising compact conformation compared to the previous low-resolution solution structure^3^, and we show that GAC can exist in both the compact and an extended conformation in solution. The overall shape of this giant armadillo-like repeat protein would enable GAC to function as an elastic linker between actin filaments and the plasma membrane and suggests a role in storing mechanical energy as myosin A (MyoA) undergoes its power stroke.

## Results

### GAC is a giant solenoid that can exist in a compact or an extended conformation

The crystal structure of FL-*Tg*GAC was determined to 2.3 Å resolution (**Table 1)**. The final model contains amino acids from Lys7 to Phe2639, missing only the first six N-terminal residues and comprising 73% α-helices, 1.4% β-strands, and 28% loop regions. FL-*Tg*GAC in the crystal has a compact solenoid structure with 164 α-helices, connected by short loops, forming 53 continuous armadillo or Huntingtin, elongation factor 3, protein phosphatase 2A (HEAT) -like repeat units (RUs)^5^, ending in the C-terminal PH domain (**Fig. 1a**). Also the loop connections between the RUs are mostly short, comprising 3-5 residues. Of note, there is a 45-Å long α-helix protruding from the end of RU 50, linking it to the first helix of RU 51 through a long loop formed by Gln2280-Ala2311. Surprisingly, this long insertion does not influence the solenoid trajectory and is only present in *T. gondii* and *Neospora caninum* (**Fig. 1** and **Extended Data Figs. 1** and **2**), which are closely related parasites in family Sarcocystidae within Apicomplexa^6^.

**Table 1.** FL-*Tg*GAC crystal structure data collection and refinement statistics.

|  | Phasing * | Native * |
| --- | --- | --- |
| **Data collection \*\*** |  |  |
| Space group | P212121 | P212121 |
| Cell dimensions |  |  |
| *a*, *b*, *c* (Å) | 121.5 148.7 185.7 | 121.4 146.7 185.9 |
| $\alpha$, $\beta$, $\gamma$ (°) | 90, 90, 90 | 90, 90, 90 |
| Resolution (Å) | 48 - 2.65 (2.815 2.65) | 47 - 2.33 (2.39 - 2.33) |
| No. of total reflections | 1319718 (202072) | 1915291 (142575) |
| No. of unique reflections | 188583 (29567) | 141753 (10315) |
| *R*<sub>merge</sub> | 0.17 (2.02) | 0.12 (3.08) |
| CC (½) (%) | 99 (41) | 100 (53) |
| *I* / $\sigma I$ | 8.9 (0.9) | 14.2 (1.2) |
| Completeness (%) | 99.4 (96.5) | 99.9 (99.5) |
| Redundancy | 7.0 (6.8) | 13.5 (13.8) |
| **Refinement \*\*** |  |  |
| Resolution (Å) |  | 47 - 2.33 (2.41 - 2.33) |
| No. of reflections used |  | 141694 (13984) |
| No. of TLS groups |  | 3 |
| *R*<sub>work</sub> / *R*<sub>free</sub> |  | 0.2296 (0.4061) / 0.2720 (0.4581) |
| No. protein residues |  | 2633 |
| No. atoms |  |  |
| Protein |  | 19955 |
| Ligand/ion |  | 9 |
| Water |  | 62 |
| Average *B*-factors |  |  |
| Protein |  | 78.69 |
| Ligand/ion |  | 101.71 |
| Water |  | 69.75 |
| R.m.s. deviations |  |  |
| Bond lengths (Å) |  | 0.002 |
| Bond angles (°) |  | 0.53 |
| Ramachandran plot |  |  |
| Favored (%) |  | 96.6 |
| Additionally allowed |  | 3.3 |
| Outliers |  | 0.1 |

\*A single crystal used for data collection. \*\*Values in parentheses are for highest-resolution shell.

**Fig 1.**
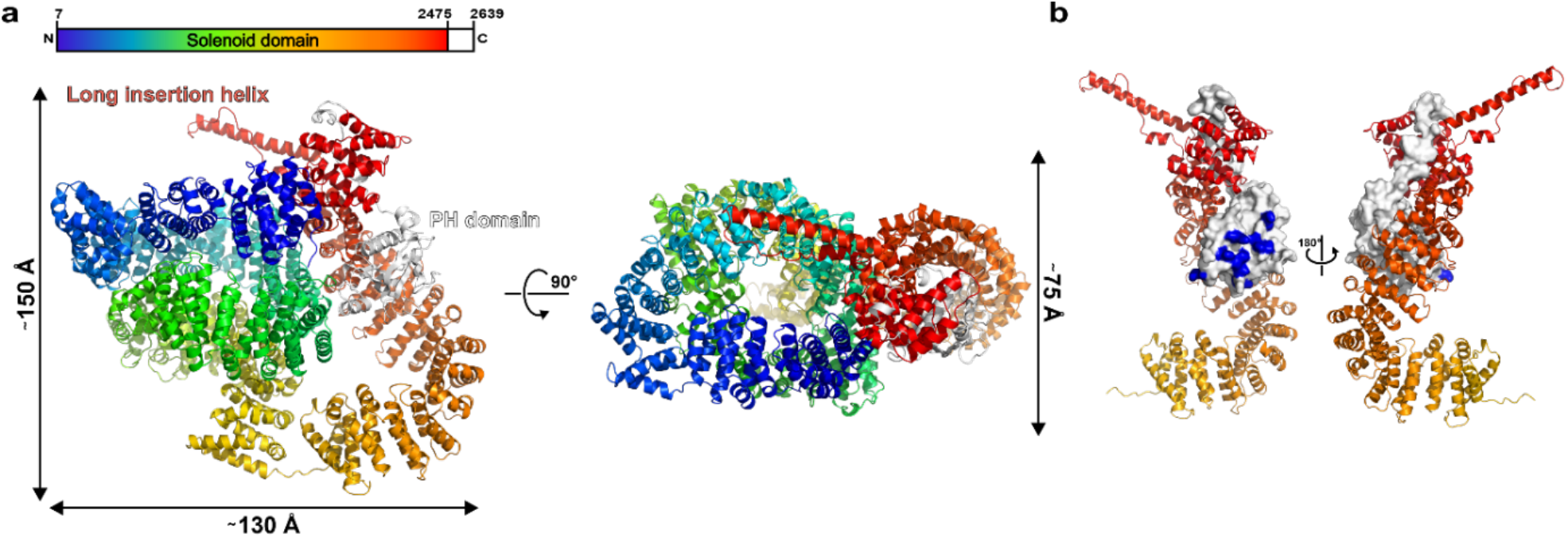
Crystal structure of FL-*Tg*GAC. a. The overall crystal structure and domain organization of *Tg*GAC. The solenoid domain from the N to the C terminus is represented in rainbow colors from blue to red, respectively. The PH domain is shown in light gray. The image on the right (“top view”) is rotated 90° along the X axis compared to the image on the left (“side view”). The solenoid domain consists of residues 7-2475 and the PH domain residues 2476-2639. The first six residues were not visible in the structure. The approximate dimensions of the structure are indicated on the side. b. The packing of the PH domain against coil 3. Coil 3 is shown as cartoon in shades of red and the PH domain surface in light gray. The Lys/Arg residues forming the non-canonical lipid-binding site are colored blue. The two images are rotated by 180° along the Y axis.

Following the solenoid domain, starting from Ser2476, is a loop region connected to the PH domain (Ser2512-Phe2639) at the C terminus of *Tg*GAC (**Fig. 1b**). Interestingly, the C-terminal region with the PH domain, which was identified as a distal adhesin-binding tip towards the plasma membrane, folds close to the N-terminal actin-filament binding region, stacking against RUs 42-51 (**Fig. 1**). This leads to the crystal structure, with approximate dimensions of 150 × 130 × 75 Å, presenting a much more compact structure in comparison to our previous low-resolution SAXS envelope, obtained from batch mode data collection^3^.

To further elucidate the configuration of FL-*Tg*GAC in solution, we prepared *Tg*GAC samples with and without crosslinking by glutaraldehyde for negative staining and cryo-EM. Native FL-*Tg*GAC appeared as elongated and heterogenous particles when viewed by negative staining or cryo-EM (**Fig. 2a**). When crosslinked, it adopted a compact shape that closely resembles the crystal structure. The crosslinked sample allowed for reconstructing a single particle cryo-EM structure of FL-*Tg*GAC at 7.6 Å (**Supplementary Data Table 1**), where well-resolved α-helices can be seen in the N-terminal solenoid region. However, slightly weaker density is observed for the C-terminal solenoid and the PH domain, suggesting flexibility of this region (**Fig. 2b** and **c** and **Extended Data Fig. 3**). In a size-exclusion chromatography (SEC) coupled SAXS experiment, FL-*Tg*GAC eluted from the column in a compact conformation, with a globular shape and a D_max_ of 152 Å, closely resembling the conformation seen in the crystal structure (**Fig. 2d, Extended Data Fig. 4a**, and **Supplementary Data Table 2**).

**Fig 2.**
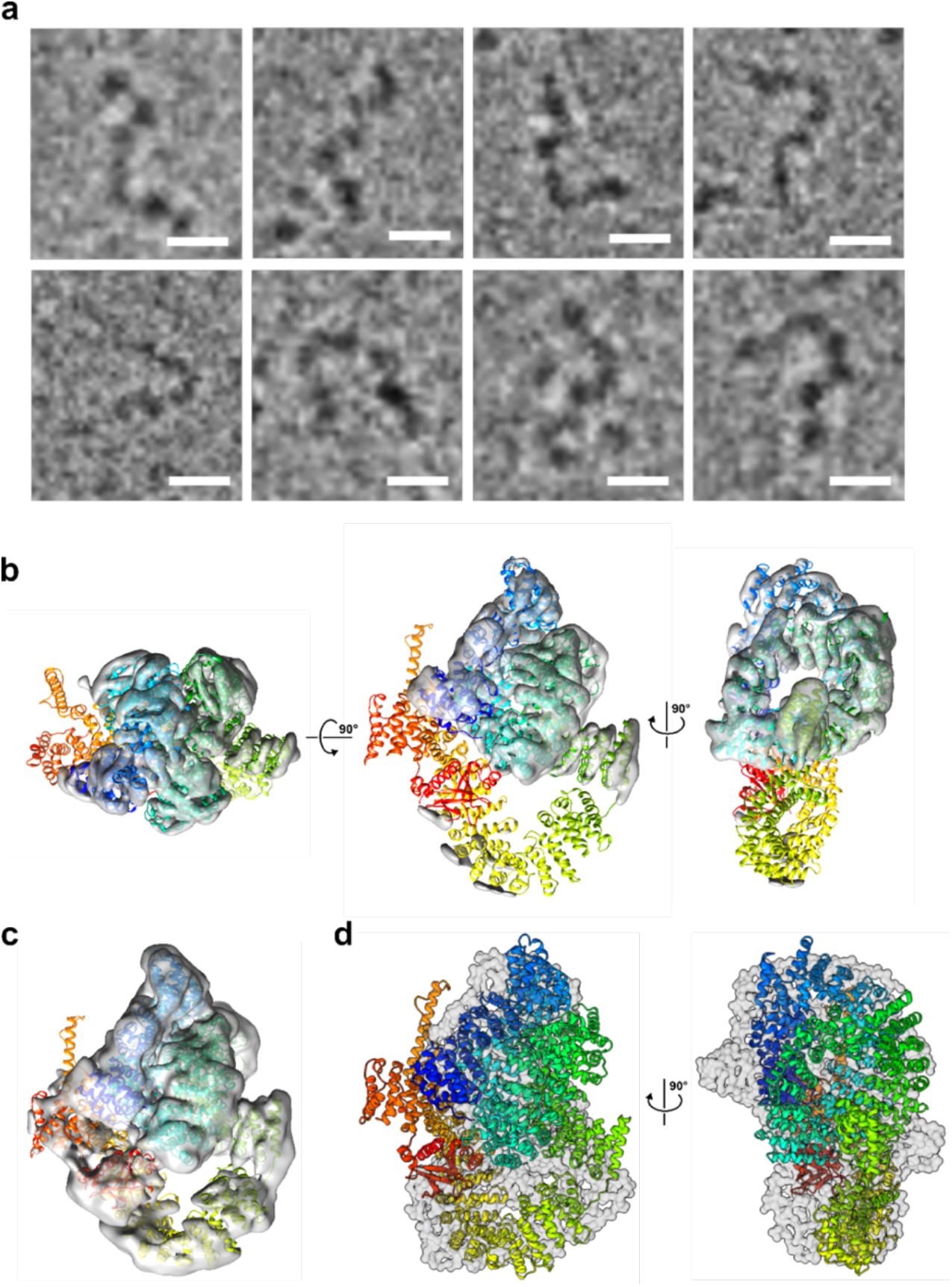
Cryo-EM results of FL-*Tg*GAC. a. Representative images of individual particles selected from the native FL-*Tg*GAC data set show the extended form and flexibility of the native sample. Scale bars represent 10 nm. b. Local filtered cryo-EM map of crosslinked *Tg*GAC with the crystal structure fit into the volume in three different orientations, 90° rotated with respect to the middle one. The threshold of the map is 2.2 σ. c. The same map as in (b) with the threshold set to 1 σ, showing the density for the whole molecule. d. Manual superposition of the FL-*Tg*GAC crystal structure onto the SEC-SAXS envelope in two different orientations, 90° apart.

### Subdomains of the GAC solenoid domain

The first and last RUs of the solenoid act as capping RUs^5,7^ with conserved hydrophobic residues facing towards the hydrophobic core to protect it from solvent and to stabilize the structure. The outer (convex) and inner (concave) helices of the RUs are amphipathic^4^ and stack approximately in parallel to the RUs with conserved Val, Ile, and Leu side chains pointing towards counterparts of neighboring repeats, contributing to a concealed hydrophobic space spanning the entire chain. Across Apicomplexa, the solvent-exposed residues are less conserved than the buried Val-Ile-Leu (VIL) clusters (**Extended Data Figs. 1** and **2**). The whole flexible solenoid domain, constituting 93% of FL-*Tg*GAC, seems to be supported by the stacking of these hydrophobic VIL clusters (**Extended Data Fig. 1**). Most of the inner helices in the RUs are tilted by approximately 15° relative to the neighboring inner helices^8^. The crossover angle between RUs is influenced by the distribution of the hydrophobic residues around the interface between RUs and the length of the linker turns/loops^9,10^. Kinks on the outer helices affect the curvature of the RU chain as well^11^. As the RUs stack with a right-handed twist, the trajectory of the RU chain twists and circulates into a right-handed solenoid^12^.

From the top view, looking down from the N-terminal part towards the C terminus, the RU chain forms three major circular coils (**Fig. 3**). Notably, between RUs 6-7, 13-14, 19-20, and 27-28, the inner helices are tilted by approximately 90° to the next inner helix, resulting in four significant kinks in the solenoid structure. The top circular coil, here named coil 1, is constituted by RUs 1 to 19 (**Fig. 3a**). After the third kink, starting from RU 20, coil 2 spirals to RU 37 (**Fig. 3b**). Of note, from RU 30 onwards, the spiral trajectory of the RU chain becomes narrower, forming a funnel-like shape (**Fig. 3b**). In contrast to coils 1 and 2, which are formed by alternating armadillo/HEAT RUs, coil 3 (RUs 38-53) is mainly composed of armadillo repeats that stack into a narrow spiral without any significant kinks (**Fig. 3c**). Coil 3 ends in a loop region before the C-terminal PH domain (**Fig. 3d**). Coils 1 and 3 contain more conserved VIL clusters than coil 2 (**Extended Data Fig. 1**), which may indicate different structural or functional stability. The interactions between coil 3 and coil 1 are a mixture of hydrophobic interactions and hydrogen bonds as well as salt bridges between residues from the N-terminal loop and RU 1 (coil 1) and RUs 49-50 (coil 3). There are fewer contacts between coil 3 and coil 2, a salt bridge between Glu2207 and Arg765 being the main interaction point. In addition, the PH domain interacts with coil 2, as discussed below.

**Fig. 3.**
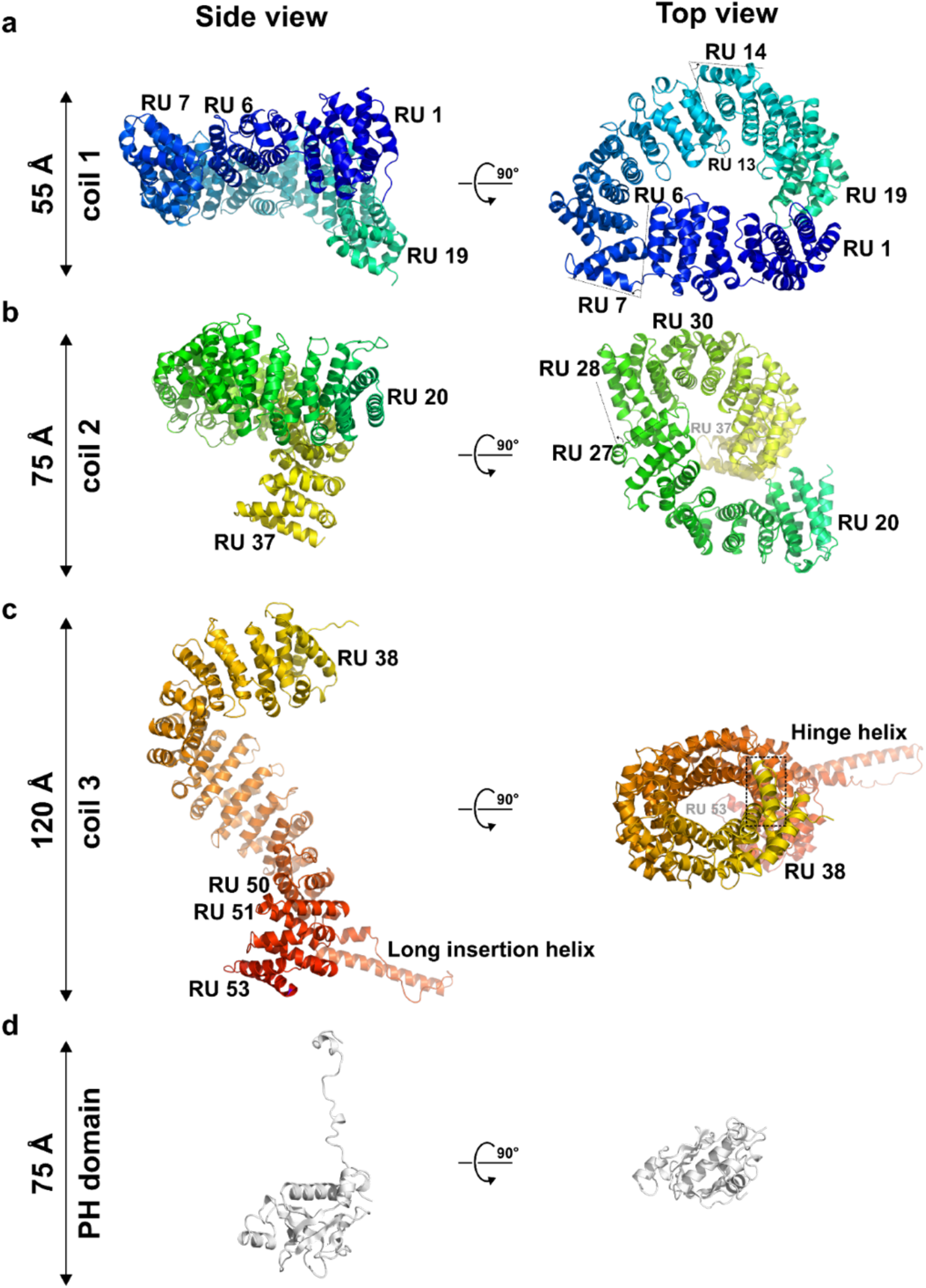
The subdomains of *Tg*GAC. For visualization purposes, the subdomains have been separated in the figure and are colored as in Fig. 1 with rainbow colors for the solenoid domain and the PH domain in light gray. The left side shows the “side view” and the right side the “top view”, which is rotated by 90° about the X axis compared to the side view. The approximate dimensions of each of the subdomains are indicated on the side. a. Coil 1 is formed by RUs 1-19. RUs 6-7 and 13-14 are hinge regions, where the solenoid domain turns and forms a spiral conformation. b. From coil 1, the solenoid domain continues down to RU 20 at the beginning of coil 2, which ends at RU 37. RUs 27-28 form a hinge region, which allows the solenoid domain to spiral downwards to coil 3 with a decreasing radius. c. Followed by the hinge helix, coil 3 is constituted by RUs 38-53, continuing the trajectory screwing downwards with no influence from the long insertion helix between RUs 50 and 51. d. The PH domain is linked to the solenoid domain with a flexible loop region, which in the crystal structure turns back, so that the PH domain sits in the groove of coil 3.

Previously, *Tg*GAC had been divided into three regions (N-terminal, middle, and C-terminal), based on sequence only^3^. Of these, only FL-*Tg*GAC and the N-terminal region (N-*Tg*GAC, residues 1-1117) could be purified in sufficient amounts for biochemical and structural characterization^3^. Based on the crystal structure, we redefined potential functional subdomain boundaries and produced new truncated forms of *Tg*GAC, hereafter called subdomains. The new constructs encoded for the full solenoid domain formed by coils 1-2-3 (*Tg*GAC coil 1-2-3; residues 1-2477), the N-terminal solenoid domain formed by coils 1-2 (*Tg*GAC coil 1-2; residues 1-1664), the C-terminal solenoid domain, *i*.*e*. coil 3 (*Tg*GAC coil 3; residues 1664-2477), and coil 3 with the PH domain (*Tg*GAC C3-PH; residues 1664-2639) (**Fig. 3, Extended Data Fig. 5**, and **Supplementary Data Table 3)**. In SEC, all other subdomain combinations, except for coil 1-2 and the PH domain, eluted earlier than FL-*Tg*GAC (**Extended Data Fig. 5**), indicating that they have larger hydrodynamic radii than FL-*Tg*GAC, despite their smaller molecular weight.

We used EM and SEC-SAXS to further explore the conformational space of the different *Tg*GAC subdomains. While native FL-*Tg*GAC without crosslinking was too heterogenous for EM reconstruction, we could study the conformation of the subdomains using negative staining (**Supplementary Data Table 4**). When analyzing *Tg*GAC coil 1-2, coherent averages were obtained for coil 2, but no density was observed for coil 1, even when crosslinked. This is evidence for flexibility of coil 1. Crosslinking, however, resulted in a more homogeneous coil 2 structure, suggesting that crosslinking reduces intrinsic flexibility of coil 2 (**Extended Data Fig. 6**). In contrast to the flexible upper coils, *Tg*GAC coil 3 has a single, seemingly rigid conformation (**Extended Data Fig. 7**). Negatively stained micrographs of crosslinked FL-*Tg*GAC coil 1-2-3 show coils 2 and 3, but not coil 1. However, coil 3 shows a range of positions relative to coil 2 (**Fig. 4a** and **b** and **Extended Data Fig. 8**). In SEC-SAXS, *Tg*GAC coil 1-2-3 shows an open conformation and a D_max_ larger than the one observed for FL-*Tg*GAC (**Extended Data Fig. 4b** and **Supplementary Data Table 1**).

**Fig 4.**
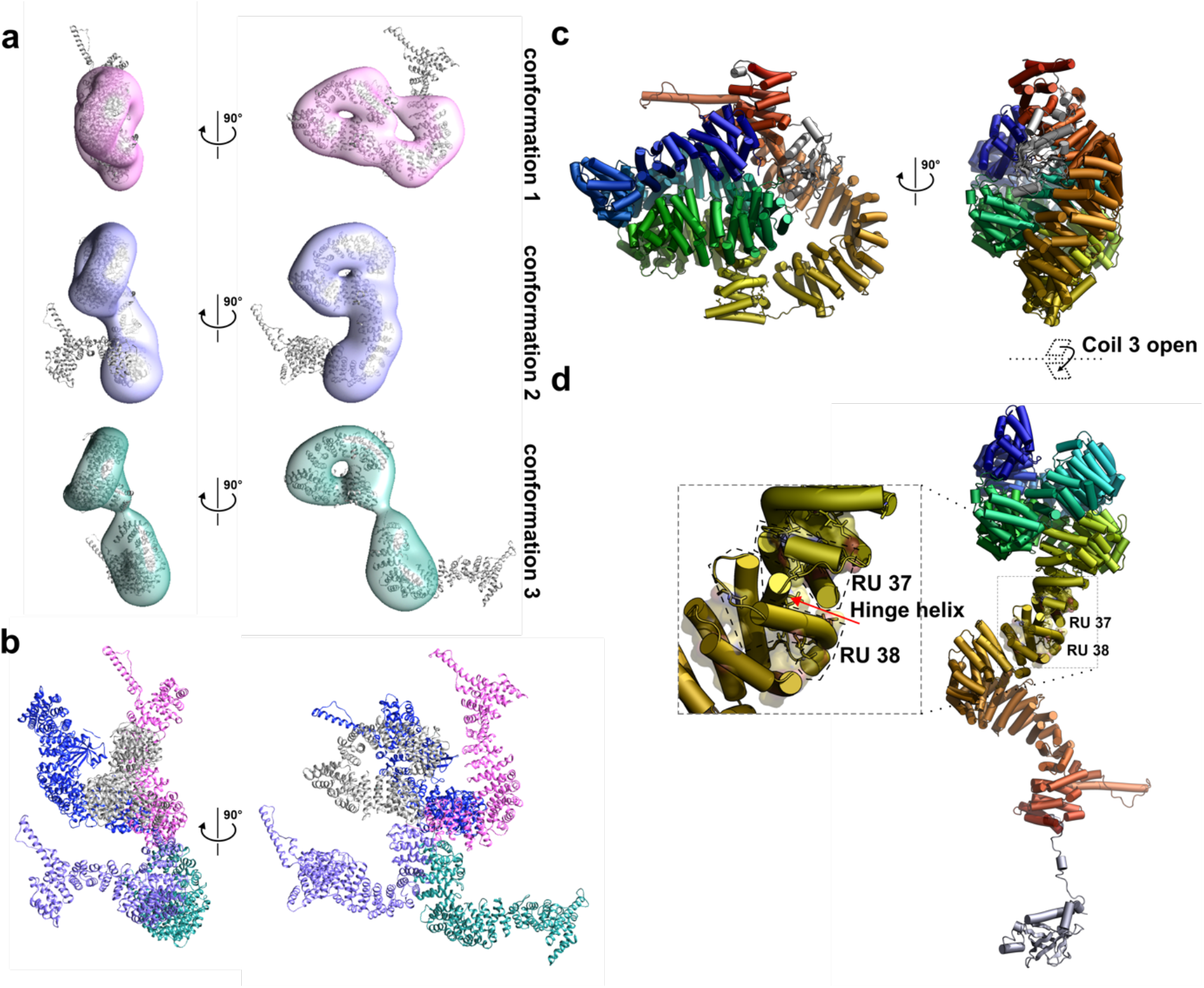
The flexible coil configuration of *Tg*GAC. a-b. Volumes obtained by 3D analysis of negative stained micrographs of crosslinked coil 1-2-3 of *Tg*GAC (no PH domain) showing three distinct ‘open’ conformations of *Tg*GAC. a. Front and top views of EM volumes corresponding to three ‘open’ conformations of *Tg*GAC with coil 2 and coil 3 models docked into the densities. b. Superposition of the three configurations in (a), with coil 2 in gray, and models of coil 3 having the same color codes as the EM maps in (a). The model in dark blue represents the position of coil 3 in the FL-*Tg*GAC crystal structure. c. The compact conformation of FL-*Tg*GAC in two different orientations, 90° apart. d. Hypothetical model of how the compact FL-*Tg*GAC in panel (c) may convert to an extended conformation upon binding to an actin filament and adhesin and/or when subject to pulling forces. The hinge region is shown in the inset, and the hinge helix is indicated with an arrow. The continuous surface of the VIL clusters would be capable of supporting a continuous super-helical conformation also from the end of coil 2 onwards. The PH domain is also shown in an opened-up conformation, but it may also stay bound to the groove of coil 3 in this model.

The combined data show that *Tg*GAC can exist in both compact and open forms, which differ in length by approximately two-fold. The open form is interpretable as a flexible association of the component coils with differing intrinsic flexibility (coil 3 < coil 2 < coil 1). The PH domain makes intimate contact with coil 3 and to a lesser extent with coil 2 (**Fig. 1** and **Extended Data Fig. 9**). Removal of the PH domain results in relative mobility of coils 2 and 3, simultaneously releasing them from coil 1 (**Fig. 4a** and **b**). This kind of mobility can be enabled by the hinge helix between RUs 37 and 38 (**Figs. 1, 3**, and **4c** and **d**). Rotating coil 3 along this hinge point would enable a continuous spiral all the way from the N-terminal coil 1 down to the end of coil 3, representing the most extended conformation possible (**Fig. 4d**).

### The PH domain has conserved membrane-binding motifs

In the crystal structure, the *Tg*GAC PH domain is snugly accommodated by the concave groove of coil 3 (**Fig. 1** and **Extended Data Fig. 9**). The core of the PH domain consists of a β-barrel formed by two perpendicular β-sheets, 4 and 5 strands each, with two strands shared between the two sheets. The C-terminal α-helix closes the barrel on one side. In addition, there are two shorter helices outside the barrel structure. Between β-strands 5 and 6, α-helix 2 of the PH domain, is amphipathic and makes contacts to coil 2 of the solenoid domain, forming a salt bridge between Lys2583 and Glu836. Also Arg2587 has several Asp/Glu residues from coil 2 at a close distance. α-helix 2 is conserved in all GACs and is present also in the PH domain of phospholipase C-δ1^13^ (PLC-δ1).

Using the FATCAT server^14^, the PH domains of PLC-δ1 (PDB ID: 1MAI)^13^ and the acylated PH (APH) domain of *T. gondii* (PDB ID: 6F8E)^11^ were identified as the closest homologs of the *Tg*GAC PH domain (**Fig. 5**). The PLC-δ1 PH domain has been studied extensively and binds stereospecifically to both PtdIns(4,5)P_2_ and Ins(1,4,5)P_3_ *via* its so-called canonical lipid binding site (**Fig. 5**)^15^. In the *Tg*GAC PH domain, this site has a reversed (negative) charge compared to both PLC-δ1 and APH PH domains (**Fig. 5**). *Tg*GAC-PH has a highly conserved Lys at position 2525 on β strand 1, followed by Phe2527 and Leu2528, which are also key residues to anchor the β1-β2 loop of APH into the membrane^11^ (**Fig. 5**). The short amphipathic α−helix is only found in GACs and the PLC-δ1 PH domain, according to a structural similarity alignment generated using iPBA^16^. In the PLC-δ1 PH domain, the corresponding α-helix is linked to the stability of IP3-binding^17,18^. In the *Tg*GAC PH domain, the last Lys at the end of this α-helix is highly conserved and a part of a KxK-like motif and could, thus, be a cooperative motif for regulation by lipid binding. Nearby, tandem KxK-Kxn(K/R)xR motifs in an extended basic patch, comprising 18 basic residues, indicate a non-canonical PA-binding surface, as seen in several other PH domains^11,19–21^. This Lys-Arg cluster is facing out towards solvent from the half-buried *Tg*GAC-PH in the groove of coil 3 and would enable lipid sensing or binding in the compact form (**Figs. 1b** and **5** and **Extended Data Fig. 9**).

**Fig. 5.**
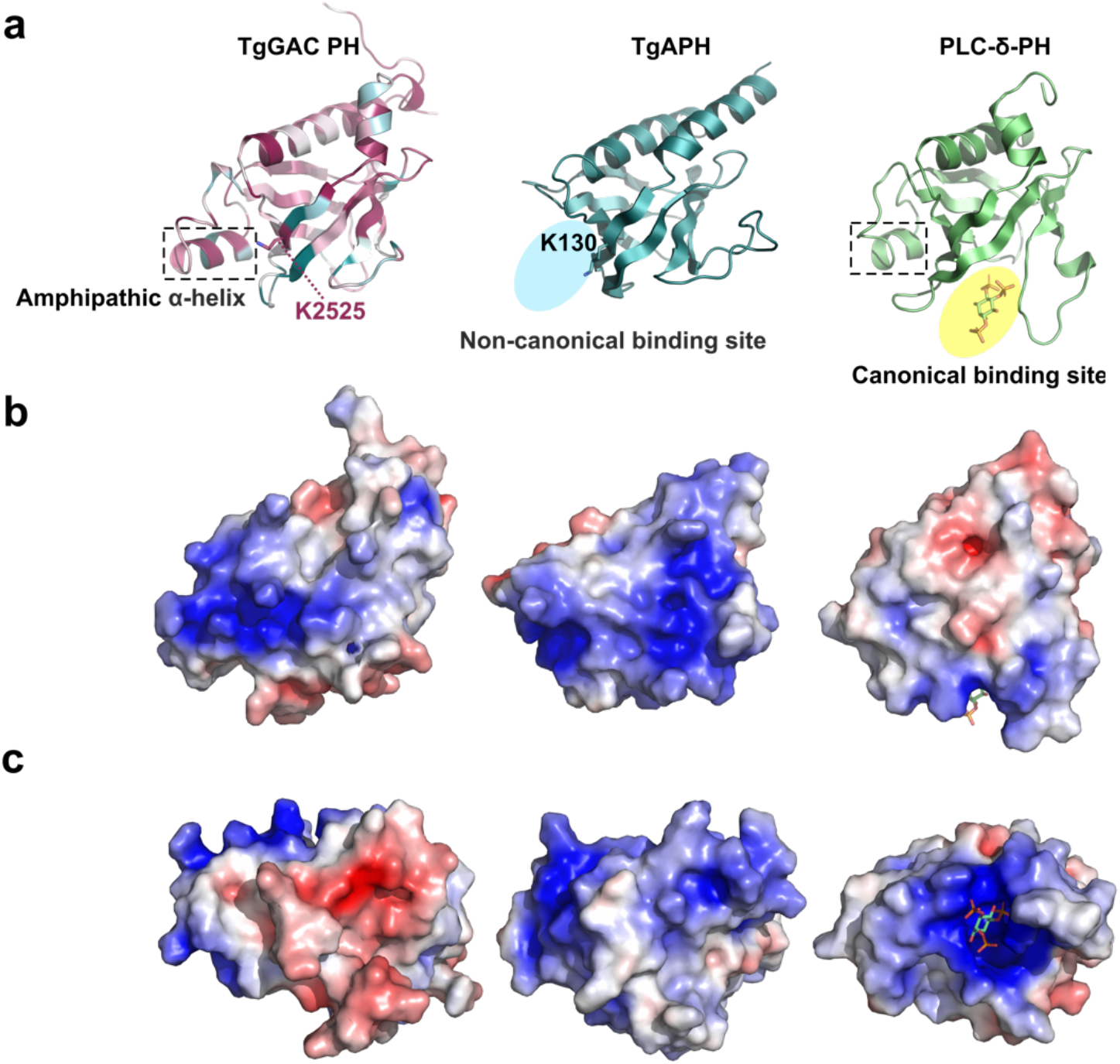
Comparison of the *Tg*GAC-PH domain with the PH domains of *Tg*APH and PLC-δ1. a. Cartoon representations of the *Tg*GAC (left), *Tg*APH (middle; PDB code 6F8E^11^), and PLC-δ1 (right; PDB code 1MAI^13^) PH domains. The canonical and non-canonical lipid binding sites are indicated in yellow and blue, respectively. The amphipathic α-helices in *Tg*GAC and PLC-δ1 PH domains are indicated with dotted line boxes, and the conserved PA-binding Lys residues in *Tg*GAC (K2525) and APH PH (K130) domains are shown as sticks and labeled. The *Tg*GAC-PH domain is colored by the ConSurf conservation score, where dark purple denotes conserved and dark cyan non-conserved. *Tg*APH is shown in cyan and PLC-δ1 in green. b. Electrostatic surface of the three PH domains in the same orientation as in panel (a). c. Electrostatic surface of the three PH domains rotated by 90° along the X axes as compared to panel (b). The bound lipid (lns(1,4,5)P_3_) in the canonical binding site of PLC-δ1 is shown as a stick model in all panels.

In summary, the *Tg*GAC PH domain contains conserved lipid-binding residues, which are exposed to solvent in the compact form. This suggests, indeed, a role in membrane recognition/interactions. A second function for the PH domain appears to be to lock coil 3 to coils 1 and 2 in the compact form.

### *The entire* Tg*GAC solenoid domain binds to actin*

Our previous results identified the N-terminal region (N-*Tg*GAC, residues 1-1117) as the cytosolic actin-filament-binding region^3^. The following middle-region (M-*Tg*GAC, residues 1118-1968) was localized apically *in vivo* and not found to interact with actin filaments *in vitro*^3^. The C-terminal region (C-*Tg*GAC, 1512-2639) was poorly expressed *in vivo* and insoluble *in vitro*^3^. With the new subdomains and the surprising positioning of the PH domain and coil 3, we decided to map more carefully the actin-binding site of *Tg*GAC using actin cosedimentation assays (**Fig. 6**). The different *Tg*GAC subdomains were all in the soluble fraction in the absence of actin. Surprisingly, all combinations containing any of *Tg*GAC coils 1 to 3 cosedimented with actin, although coil 1-2-3 seemed to bind to actin in a higher ratio than FL-*Tg*GAC. All the *Tg*GAC coils cosedimented with both *Pf*ActI, which is a close homolog of *Tg*Act, and vertebrate skeletal muscle α-actin filaments but seemed to have higher affinity to *Pf*ActI than to α-actin (**Fig. 6**). The PH domain (residues 2501-2639) did not interact with either *Pf*ActI or α-actin. Thus, the whole solenoid domain seems to be involved in or is at least capable of actin binding. The variable conformations adapted by FL-*Tg*GAC in solution and the attachment of the PH domain to coil 1 in the closed conformation seem to affect the ability of *Tg*GAC to bind to actin.

**Fig 6.**
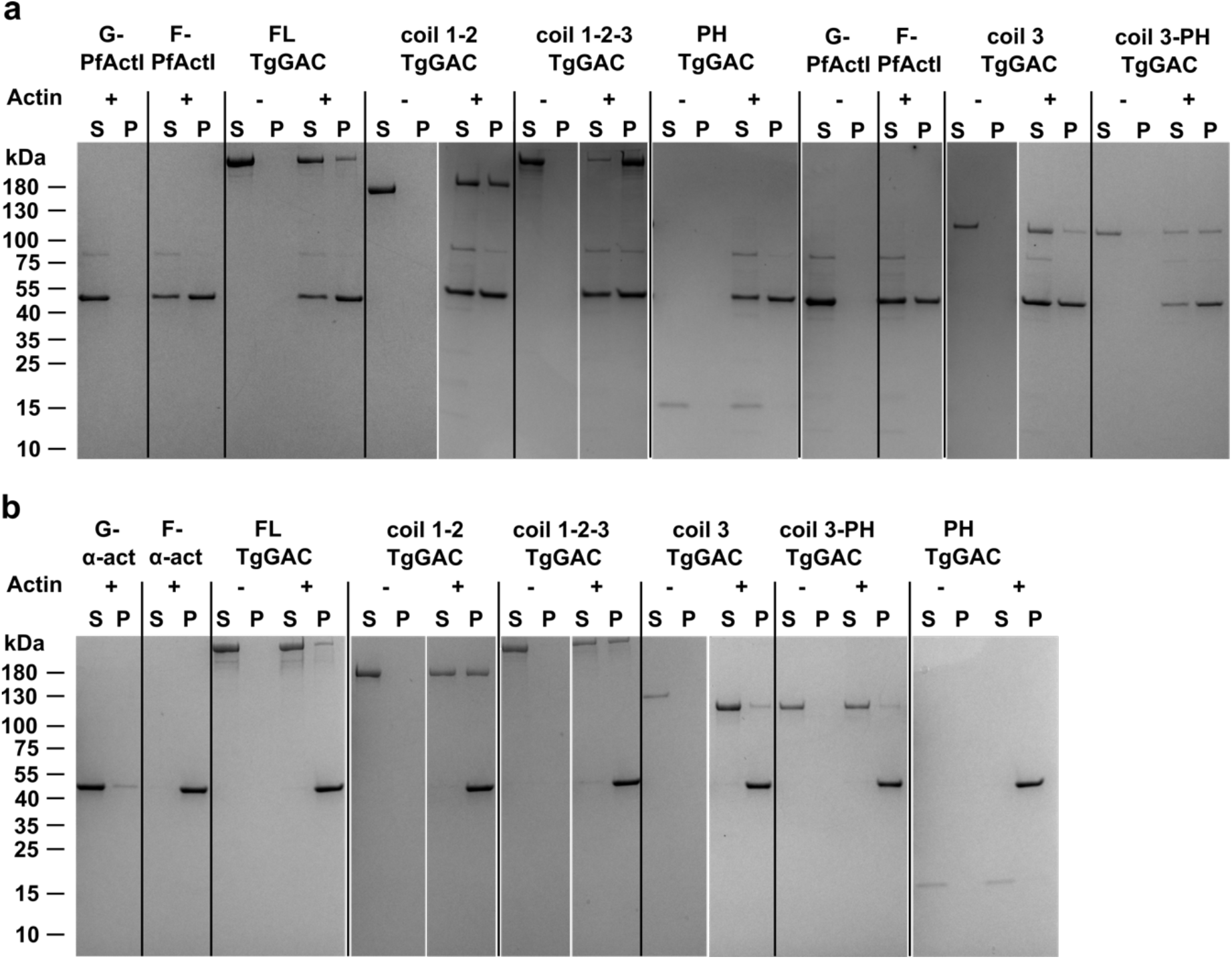
Cosedimentation of *Tg*GAC with actin. a. FL-*Tg*GAC and all the subdomains, except for the PH domain, cosedimented with filamentous *Pf*ActI. The experiment was performed as biological triplicates and shown are representative gels of three. b. FL-*Tg*GAC and all the subdomains, except for the PH domain, cosedimented with vertebrate skeletal muscle α-actin. This control experiment was performed only once. In both panels, S denotes the supernatant and P the pellet fractions. G-actin served as an actin quality control. The molecular weight marker sizes in kDa are indicated to the left of the gels.

## Discussion

### Functional insight into the solenoid domain

The *Tg*GAC solenoid domain makes three helical turns with a radius that progressively narrows, moving from the N terminus towards the C-terminal, membrane-binding, PH domain. In a partially extended form, it would resemble a conical compression spring bridging between actin and membrane attachment sites. In the compact form, the structure is twisted upon itself by at least two hinge regions. This compact structure also exists in solution but has the tendency to extend under certain conditions or over time, as seen from our previous batch SAXS experiments^3^ and the native EM structures. The solenoid spiral architecture is enabled and maintained by the arrangement of the armadillo/HEAT-like RUs and the conserved hydrophobic core, formed by the VIL clusters. Such a hydrophobic core can be a significant factor driving protein folding to maintain structure and function^4^. In the *Tg*GAC solenoid domain, the hydrophobic core extends throughout the whole RU chain, maintaining not only local folding of each RU but also retaining the tertiary structure. The solenoid domain might therefore undergo significant deformation without disrupting secondary or tertiary structures^22,23^. Considering the placement of GAC in the glideosome complex, and its potential to behave as an extensible, spring-like, connection between actin and the plasma membrane it might be expected to play a mechanical functional role.

While we previously mapped actin binding to the N-terminal coil region, we now, with the newly designed subdomains, show that the entire solenoid domain interacts with actin, and only the C-terminal PH domain does not. Based on cosedimentation assays, all combinations containing coils 1 and 2 but not the PH domain bind actin more effectively than the FL-*Tg*GAC or coil 3. The most conserved parts, in addition to the VIL clusters throughout the solenoid, are found in coil 3 and the top surface of coil 1 (**Extended Data Figs. 1** and **2**), which may indicate the most important regions. The conservation of coil 3 may reflect the importance of its interactions with the PH domain, and the more efficient actin binding in the absence of the PH domain supports a regulatory function of the PH-coil 3 interaction.

Since the entire solenoid binds actin this means GAC might lie parallel between the actin filament and the plasma membrane consistent with the limited space of approximately 30 nm between the plasma membrane and the IMC, of which the actomyosin complex already occupies approximately half. This lateral binding mode would also be compatible with stabilization of actin filaments. However, it is not known whether all the coils bind to actin simultaneously or if sequential binding events take place as MyoA moves actin rearwards. It seems plausible that parts of the binding interface may only be exposed as the compact structure extends, given the higher binding ratio in the absence of the PH domain that locks the FL protein in the compact conformation.

### Role of the PH domain

A prerequisite for gliding motility and invasion is that adhesin molecules, present on the outer surface of the parasite, must anchor to the host cell surface or the moving junction^24,25^. *Tg*GAC is immobilized to the inner leaflet of the parasite plasma membrane by binding to the intra-cellular portion of adhesin and PA in the membrane *via* its C-terminal solenoid domain and/or the PH domain. PH domains are versatile modules for protein-protein and protein-membrane interactions, mostly in eukaryotic cells^21^. The PH domain of *Tg*GAC is quite distant even from its closest homologs, PLC-δ1 and the APH domain-containing protein. Although sequence identities are very low, overall structures are conserved. Structurally, the *Tg*GAC PH domain more closely resembles PLC-δ1 but functionally, it seems closer to the APH PH domain with the extended non-canonical lipid binding site and PA binding. The APH domain-containing protein is essential for motility, cell entry, and egress of *P. falciparum* and *T. gondii*. It is recruited to PA-enriched membranes *via* both canonical and non-canonical lipid binding sites (**Fig. 5**)^11^.

We have previously shown that *Tg*GAC transiently associates with *Tg*MIC2 and that the *Tg*GAC PH domain binds PA and adhesin tails^3^. This would be in line with a location of the PH domain at the periphery of FL-*Tg*GAC, far from the actin-binding region, as proposed in our previous extended SAXS model^3^. The closed conformation stabilized by the PH domain may be important for recruitment to the conoid at the apical tip, which is also dependent on methylation by the apical complex lysine (K) methyltransferase (AKMT)^3,26^. It is possible that GAC would then, upon binding to both an actin filament and the plasma membrane, adopt an extended structure. Thus, PH domain membrane interactions may function as a switch, leading to the compact GAC structure extending upon PA-enriched membrane and actin binding (**Fig. 4c** and **d**).

### Elastic spring model for GAC in gliding motility

Given the elastic characteristics of solenoid domains^22,23,27–36^ and examples of helical repeat proteins, like the catenin core complex in adherens junctions^37,38^ and talin in focal adhesions^39,40^, it seems plausible that GAC may act as a spring-like connection between the short (∼100 nm) actin filaments^41–43^ and the adhesion complex that is formed during gliding motility and invasion (**Fig. 7**). This could be analogous to the ankyrin repeats in the NOMPC-TRP ion channel, which have been suggested to play an important role in its mechanosensory function ^44^. GAC may cluster or accumulate during gliding and invasion, as suggested by the observation of intense ring-like structures at the moving junction^3^. GAC molecules presumably accumulate along with the apico-basal waves of filamentous actin and adhesins^3,45^ to sustain mechanical force during gliding.

**Fig 7.**
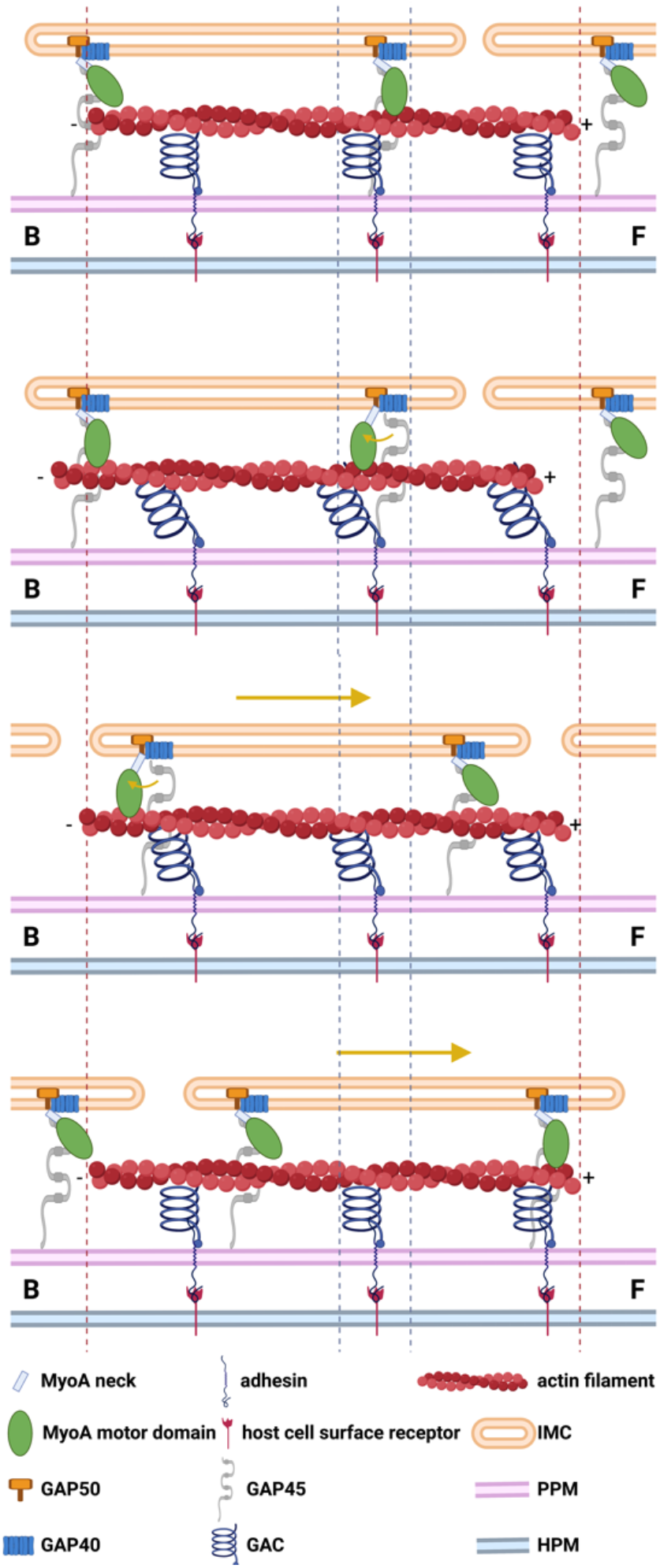
The spring model of GAC as an elastic element in the glideosome. a. GAC is in a relaxed (contracted) conformation, conneting the actin filament to the adhesin in the parasite plasma membrane (PPM). b. The MyoA power stroke shifts back the actin filament, leading to extension of the GAC solenoid structure. Partial (c) and full (d) relaxation of the solenoid domain to the contracted state lead to forward movement of the actin filament and the IMC relative to the adhesin in the PPM and host plasma membrane (HPM). The GAC-actin filament units can be periodically powered by successive myosins to transport the IMC-myosin complex toward the apical end and eventually move the whole parasite forward. GAC as an elastic element allows each actin filament to act as an independent unit, releasing the requirement of synchronization of all the myosins. Depicted are GAC, an actin filament, MyoA motor and neck domains, gliding associated proteins (GAP) 50, 40, and 45, an adhesin, a host cell surface receptor, the IMC, PPM, and HPM. The large arrows represent the forward movement of the parasite, the small arrows represent the MyoA power stroke direction. F denotes the apical (front) and B the posterior (back) end of the parasite. The figure was generated using BioRender.

Based on the localization of GAC and its hinged conical spring-like architecture, we propose a role for GAC in storing and transmitting mechanical work and force during gliding motility and host cell invasion (**Fig. 7**). Force from the MyoA power-stroke is transferred in series, across the actin-GAC complex, to the adhesin, driving the adhesion site rearwards, towards the basal end of the parasite. GAC might act as an elastic storage element, allowing the force generated by several MyoA molecules to sum, thereby producing sufficient force to drive the parasite into the red blood cell during invasion. Relaxation of the extended GAC spring would maintain the force required to drive the IMC forwards relative to the rearward (basal) movement of the tight-junction region that is observed as the merozoite penetrates the red blood cell.

#### Concluding remarks

GAC is a protein unique to and conserved across apicomplexan parasites and indispensable for their gliding motility. It is the largest armadillo-repeat protein in Apicomplexa^4^ and the largest armadillo-repeat protein, for which a structure is known to date. GAC has two-fold conformational flexibility: (i) The C-terminal PH domain anchors it in the compact conformation and may function as a sensor for lipid binding and as a switch between the open and closed states. (ii) The solenoid domain confers spring-like flexibility storing mechanical energy for gliding motility.

## Methods

### *Expression and purification of FL-*Tg*GAC and the subdomains*

The FL-*Tg*GAC construct was modified from the His-FL-*Tg*GAC published before^3^ to add a GST tag at the N terminus, followed by a TEV protease site, and an uncleavable 6×His tag at the C terminus. All the subdomains (**Supplementary Data Table 2**) were constructed using the same fusion tag strategy.

Selenomethionine-derivatized FL-*Tg*GAC was expressed in the methionine auxotroph T7 Express Crystal *Escherichia coli* strain (NEB #C3022) cultured in SelenoMethionine Medium Base plus Nutrient Mix (Molecular Dimensions MD12-501), supplemented with 25 mg l^-1^ kanamycin and 40 mg l^-1^ L(+)-Selenomethionine (Acros Organics). The culture was incubated at 37°C while shaking at 200 r.p.m. until the optical density at 600 nm (OD600) reached 1.0. FL-*Tg*GAC expression was then induced with 0.75 mM isopropyl-β-D-thiogalactoside (IPTG) and the culture continued at 18°C for 20 h. Native FL-*Tg*GAC and all the subdomains were produced in *E. coli* BL21 (DE3) using auto-induction ZY-5052 medium supplemented with 25 mg l^-1^ kanamycin at 18°C for 24 h.

All the native and Se-Met-derivatized *Tg*GAC proteins were purified from sonicated cell lysates in buffer A (50 mM Tris-HCl, pH 7.5, 500 mM NaCl) with 10 mM TCEP, 0.2 mg ml^-1^ lysozyme, and SIGMAFAST™ Protease Inhibitor Cocktail Tablets. The lysate was clarified by centrifugation at 18000 g for 35 min at 4°C and then incubated with Pierce™ Glutathione Agarose (Thermo Scientific™) equilibrated with buffer A with 1 mM TCEP at 10°C for 1 h. Following the incubation, the protein-agarose mixture was poured and washed in a gravity-flow column with buffer A with 1 mM TCEP and then buffer B (20 mM Tris-HCl, pH 7.5, 50 mM NaCl, 0.5 mM TCEP) for 5 column volumes (CV), each. The bound FL-*Tg*GAC was eluted after incubation with TEV protease in buffer B and subsequently loaded onto a HisPur™ Ni-NTA Resin column (Thermo Scientific™). Then, 5 CV of 15 mM and 80 mM imidazole containing buffer B were applied to wash the resin and to elute the FL-*Tg*GAC, respectively. The purified FL-*Tg*GAC was concentrated and applied to a size exclusion column (Superdex 200 Increase, GE Healthcare), equilibrated with buffer C (10 mM Tris-HCl, pH 7.5, 50 mM ammonium acetate, 0.2 mM TCEP). The final purified FL-*Tg*GAC was concentrated to 15-20 mg ml^-1^ for crystallization.

The *Tg*GAC subdomains were expressed in *E. coli* BL21 (DE3) using auto-induction ZY-5052 medium supplemented with 25 mg l^-1^ kanamycin at 18°C for 24 h. The proteins were purified from *E. coli* lysates after sonication in buffer A supplemented with 1 mM TCEP, 0.2 mg ml^-1^ lysozyme, and SIGMAFAST™ Protease Inhibitor Cocktail Tablets. The clarified lysates were incubated with Pierce™ Glutathione Agarose (Thermo Scientific™) equilibrated with buffer A with 1 mM TCEP at 10°C for 1 h. Following the incubation, the protein-agarose mixture was poured and washed in a gravity-flow column with buffer A with 1 mM TCEP and then buffer B for 5 CV, each. The bound proteins were eluted from the beads after incubation with TEV protease in buffer B. The purified subdomains were subsequently concentrated and applied to a size exclusion column (Superdex 75 increase, GE Healthcare for *Tg*GAC-PH and Superdex 200 Increase, GE Healthcare for the other subdomains), equilibrated with buffer D (50 mM Tris-HCl, pH 7.5, 50 mM NaCl, 0.1 mM TCEP).

### Crystallization, data collection, and structure determination

Purified native and Se-Met FL-*Tg*GAC were crystallized using the hanging-drop vapor diffusion method at 16°C. Crystals grew from a 2:1 mixing ratio of FL-*Tg*GAC to reservoir solution, containing 0.1 M Tris, pH 8.4, 1.4 M potassium sodium tartrate, 50 mM magnesium acetate. All crystals were cryoprotected with cryoprotectant solution containing 0.1 M Tris, pH 8.4, 2.3 M potassium sodium tartrate, 50 mM magnesium acetate, 50 mM ammonium acetate, and 0.2 mM TCEP.

The crystal structure of FL-*Tg*GAC was determined using selenomethionine (Se-Met)-labeling and single-wavelength anomalous diffraction (SAD). The final X-ray diffraction experiments were carried out at 100 K on the protein crystallography beamline X06DA-PXIII at the Swiss Light Source, Paul Scherrer Institute, Villigen, Switzerland. SAD data of the SeMet-derivatized crystal were collected using a 50 × 90 μm^2^ X-ray beam with a wavelength of 0.9729 Å. The substructure and the preliminary model were solved from a Se-Met-labeled FL-*Tg*GAC crystal diffracting to 2.6 Å. The final high-resolution model was refined using data from a different Se-Met-FL-*Tg*GAC crystal, diffracting to 2.3 Å (**Table 1**). The SAD and native data were processed and scaled using the XDS^46^ package. HKL2MAP^47^ was used for phase determination, and CRANK2^48^ was used to build an initial model. Interactive manual model building in COOT^49^ and refinement with PHENIX.REFINE^50^, using TLS in the final rounds, against the higher resolution data were then performed. The final structure has 96.6% of residues in the favored and 3.3% in allowed regions of the Ramachandran plot (**Table 1**). The secondary structure contents were analyzed using PDBsum^51^ and the conservation using ConSurf^52^. Figures were prepared using PyMOL^53^ and ChimeraX 1.1^54^.

### *Cryo-EM sample preparation, imaging, and data processing for FL-*Tg*GAC*

FL-*Tg*GAC was diluted to 0.2 mg ml^-1^ in 20 mM HEPES, pH 8.0, 50 mM NaCl, 10 mM MgCl_2_ with or without 1 wt% glutaraldehyde (GA). The sample with GA was kept on ice for 5 min for crosslinking reaction before plunge freezing. 4 μl of the crosslinked or native sample were applied to amylamine glow-discharged (25 mA, 30 s) Quantifoil R2/2 grids in the environmental chamber of a Vitrobot Mark IV (FEI/Thermo) at 4°C and 95% humidity, and the grid was blotted with filter paper for 4 s and then plunge frozen in liquid ethane at liquid nitrogen temperature.

The grids were screened on a Talos Arctica microscope (FEI/Thermo) at 200 kV, and one of the crosslinked *Tg*GAC grids was selected for an overnight data collection using EPU v 2.11. A total of 952 ten-frame movies were collected using a Falcon III camera in linear mode, with dose rate and exposure time of 46 e/Å^2^/s and 1.52 s, respectively. The pixel size of the movies is 1.61 Å, and the defocus range was set from -1.5 to -3.5 μm with 0.5 μm interval.

Pre-processing of the movies was conducted in Relion v. 4.0^55^. The frames were aligned and dose weighted with Relion’s own motion correction (a typical micrograph shown in **Extended Data Fig. 3b**), followed by CTF estimation with CTFFIND v. 4.4.13^56^. Ten micrographs with various defocus values were selected and picked with crYOLO^57^ general model, after which refinement for the picking was done manually. The selected coordinates were then used to train a model in crYOLO v 1.8.2, which was then used to predict the coordinates of the particles from the whole dataset. In total, 569554 raw particles were picked, extracted binned 2 (pixel size 3.22 Å, box size 100), and transferred to CryoSPARC v. 3.3.1^58^ for 2D classification. After 2D classification, 136937 particles were selected and re-extracted in Relion with full size (pixel size 1.61 Å, box size 200). 3D classification was performed with ab-initial reconstruction in CryoSPARC, and 69523 particles were selected and subjected to 3D refinement (non-uniform refinement). The refined map was local filtered by Local Anisotropic Sharpening in Phenix v. 1.20.1^59^. The workflow from movies to final map is displayed in **Extended Data Fig. 3c**. The final refined map has a global resolution of 7.6 Å, calculated by gold-standard Fourier shell correlation (**Extended Data Fig. 3d**). The crystal structure of FL-*Tg*GAC was rigidly docked into the cryo-EM map in Chimera v. 1.13.1^54^ using the ‘fit in map’ function.

### *Negative stain sample preparation, imaging, and data processing for* Tg*GAC subdomains*

Structures of the *Tg*GAC subdomains were explored using the negative stain EM imaging method. The coil 3 sample was diluted to 0.2 mg ml^-1^ with the same buffer as used for FL-*Tg*GAC. The coil 1-2-3 sample was diluted to 0.2 mg ml^-1^ in the same buffer supplemented with 1 wt% GA. Coil 1-2 was diluted to 0.2 mg ml^-1^ and split into two samples with 1 wt% GA added into one of them. The samples were kept on ice for 4 min before pipetting 4 μl onto a copper mesh grid with a continuous carbon film. The sample was blotted away with filter paper from the side of the grid after 1 min, and 4 μl of 1 wt% sodium silicotungstate were immediately applied to the grid. The stain was kept on the grid for 45 s before being blotted away using the same method as above. The grid was then dried on filter paper before inserting into the microscope.

Images were acquired with a CCD camera in a Tecnai Spirit TEM (FEI) operated at 120 kV using SerialEM (v 3.7.6) for automatic data collection. For each sample, 200-300 single-frame micrographs with an exposure time of 1 s were collected. Typical images of coil 1-2 (native and crosslinked), coil 3 and coil 1-2-3 are shown in **Extended Data Fig. 6a, Extended Data Fig. 6g, Extended Data Fig. 7a**, and **Extended Data Fig. 8a**. The pixel size used was 3.3 Å, and the defocus range was -1.5 to -3 μm. The data processing protocol was identical to the cryo-EM dataset except that no motion correction was performed. The resolutions for the final refined maps are in the range of 20-23 Å. The workflow and the important steps of the data processing for the negative stain dataset are shown in **Extended Data Fig. 6**, E**xtended Data Fig. 7**, and **Extended Data Fig. 8**. The crystal structures of the corresponding fractions of *Tg*GAC were rigidly docked into the 3D maps in Chimera v. 1.13.16.

### Size exclusion chromatography-coupled small-angle X-ray scattering

SEC-SAXS data were collected on the SWING beamline^60^ at the Synchrotron SOLEIL, Paris, France. SAXS data were obtained from FL-*Tg*GAC and *Tg*GAC coil 1-2-3 at concentrations 1 and 5 mg ml^-1^, respectively. The protein samples were loaded onto an Agilent Biosec3-300 in 20 mM Tris-HCl, pH 7.5, 50 mM NaCl, 1% sucrose, 0.5 mM TCEP at 0.3 ml min^-1^. The SAXS data were recorded using a detector distance of 2 m, and exposure time of 990 ms/frame, and a wavelength of 1.033 Å. The data were processed and analyzed using programs from the ATSAS package^61^. *Ab initio* models were generated using either DAMMIN^62^ or GASBOR^63^. The crystal structure of FL-*Tg*GAC was fitted to the experimental data using CRYSOL^64^. The figures were generated using ChimeraX 1.1^54^.

### Actin cosedimentation assays

To assess the binding of the *Tg*GAC subdomains to α-actin (produced and purified as previously^65^) and *Pf*ActI (expressed and purified as previously^66^), a cosedimentation assays were performed. *Pf*ActI in G buffer (10 mM HEPES, pH 7.5, 0.2 mM CaCl_2_, 0.5 mM ATP, and 0.5 mM TCEP) and α-actin (10 mM Tris-HCl, pH 7.5, 0.2 mM CaCl_2_, 0.5 mM ATP, and 0.5 mM TCEP) were polymerized for 2 h at room temperature by adding F buffer to final concentrations of 50 mM KCl, 4 mM MgCl2, and 1 mM EGTA. The polymerized actins were mixed with the *Tg*GAC subdomains to final concentrations of 4 and 0.5 μM, respectively. The filaments were sedimented at 435000 g for 1 h at room temperature. Both pellet and supernatant fractions were analyzed using SDS-PAGE and Coomassie Brilliant Blue staining.

## Data availability

The crystal structure coordinates and structure factor amplitudes have been deposited to the Protein Data Bank (PDB code 8BT6) and the EM maps to the Electron Microscopy Data Base (EMDB codes EMD-16257, EMD-16258, EMD-16259, and EMD-16260).

## Acknowledgments

We are grateful to Dr. Justin Molloy for enlightening discussions on the manuscript, motor proteins, and the spring model and to Dr. Petri Kursula for discussions on the GAC structure and for critical reading of the manuscript. We thank Drs. Chia-Ying Huang, Vincent Olieric, and Sylvain Engilberge from Paul Scherrer Institute for valuable help during numerous data collections for obtaining the phases for the crystal structure and Aleksi Sutinen for help with SAXS data collection and preliminary data processing. We thank Dr. Andrea Nans of the Francis Crick Institute Structural Biology Science Technology Platform for advice on data collection and computing as well as Drs. Philip Walker and Andrew Purkiss of the Francis Crick Institute Structural Biology Science Technology Platform and the Francis Crick Institute Scientific Computing Science Technology Platform for computational support. We also gratefully acknowledge the following synchrotron beam lines for crystal screening as well as X-ray diffraction and SAXS data collection time: X06DA-PXIII at Swiss Light Source; P11 at DESY; P12, P13, and P14 at EMBL/DESY: SWING at Synchrotron SOLEIL; and I03 and I04 at Diamond Light Source. We thank the Academy of Finland, Sigrid Jusélius foundation, Emil Aaltonen foundation, Jane and Aatos Erkko foundation, and the Norwegian Research Council for financial support. P.B.R. was supported by the Francis Crick Institute, which receives its core funding from Cancer Research UK (CC2106), the UK Medical Research Council (CC2106), and the Wellcome Trust (CC2106). For the purpose of Open Access, the authors have applied a CC BY public copyright license to any Author Accepted Manuscript version arising from this submission.

## EXTENDED DATA FILES

**Extended Data Fig. 1.**
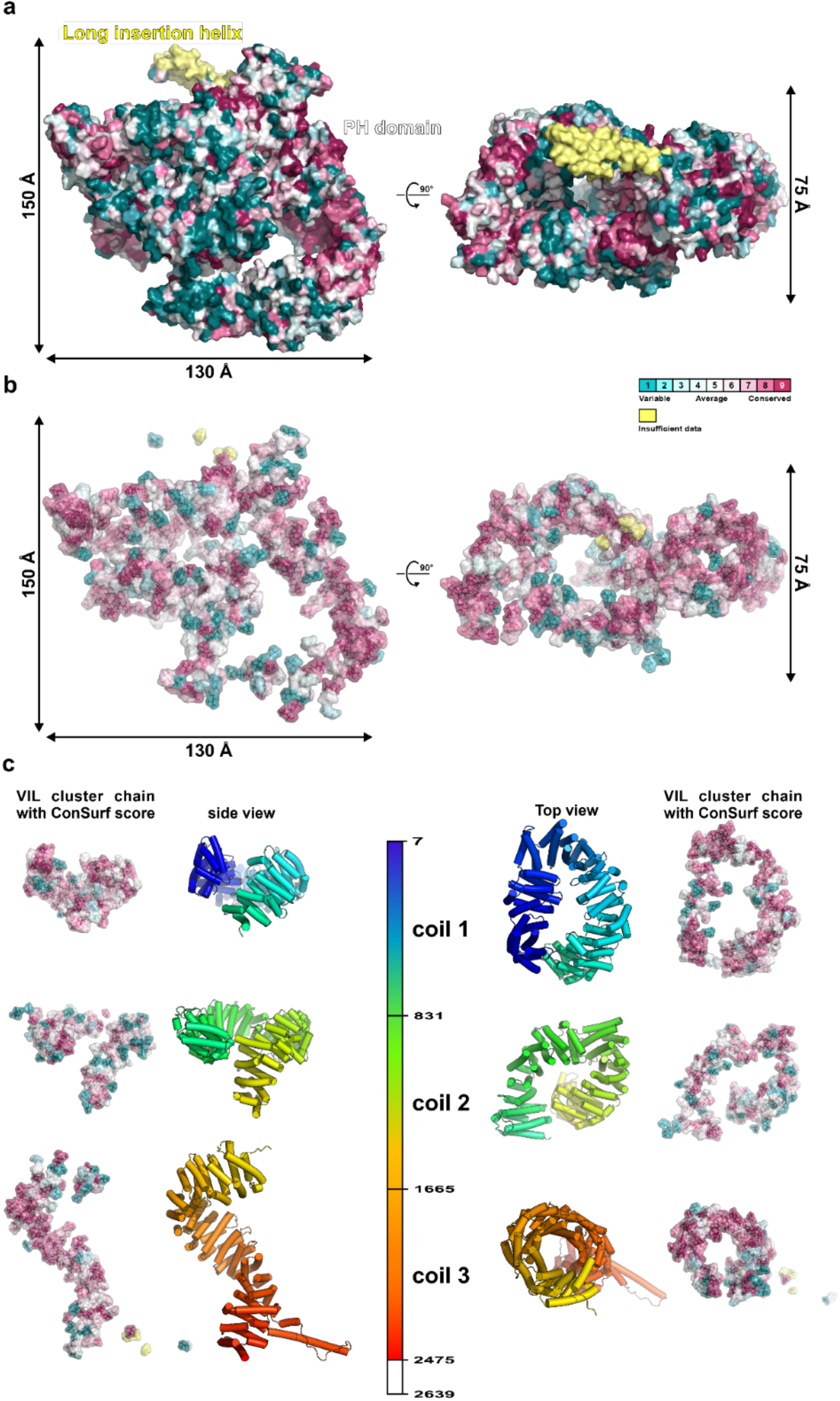
Conservation along the *Tg*GAC solenoid subdomain surface and the VIL clusters. a. The FL-*Tg*GAC surface colored by the conservation scores from ConSurf. The long insertion helix is shown in yellow due to insufficient data for conservation comparison. The C-terminal region (coil 3) of GAC possesses the largest continuous conserved surface. The N-terminal coil 1 contains some conserved regions, in particular on the “top” of the coil. The middle region contains the lowest amount of conserved VIL clusters and surface. b. The VIL cluster surface of *Tg*GAC with the same color definition as in (a), showing that the VIL clusters are generally conserved throughout the solenoid domain. c. The conservation of the VIL clusters displayed in the “opened-up” form.

**Extended Data Fig. 2.**
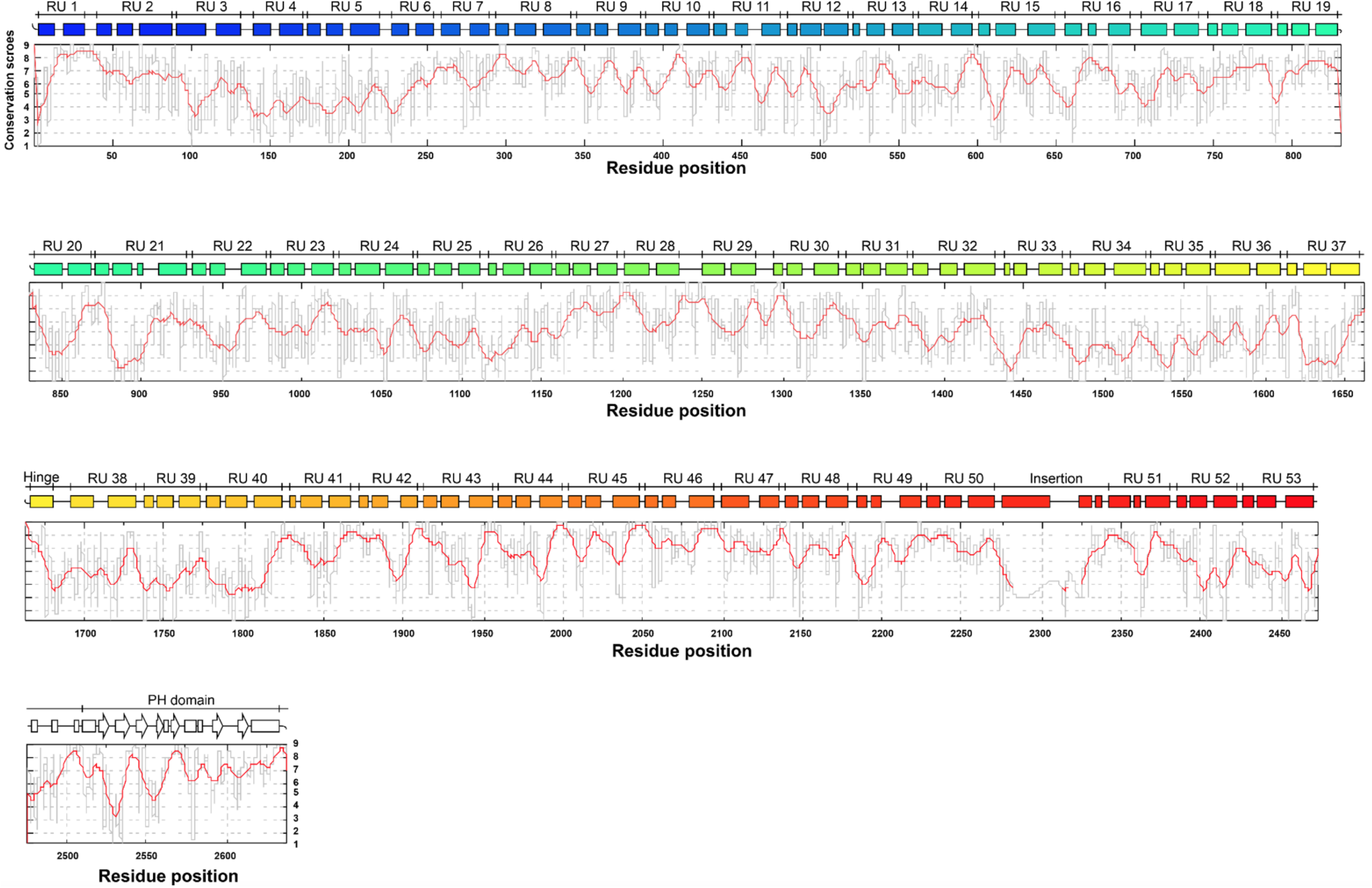
Conservation plot of the *Tg*GAC sub-solenoid domains and the organization of the RUs. The conservation score is plotted as a function of *Tg*GAC residue position across GAC sequences from apicomplexan species: *T. gondii, P. falciparum, N. caninum* Liverpool, *Piliocolobus tephrosceles, Eimeria maxima, Cyclospora cayetanensis, Besnoitia besnoiti, Babesia bigemina*, and *Theileria equi* strain WA. The grey trace indicates scores for individual residues, the red trace indicates scores averaged over an 11-residue stretch, smoothed by Origin. The secondary structure elements above the plot are colored in rainbow colors from blue to red for the solenoid domain and white for the PH domain. The RUs are numbered from 1 to 53, except for two helices, the hinge helix in between RU 37 and RU 38 and the insertion helix in between RU 50 and 51.

**Extended Data Fig. 3.**
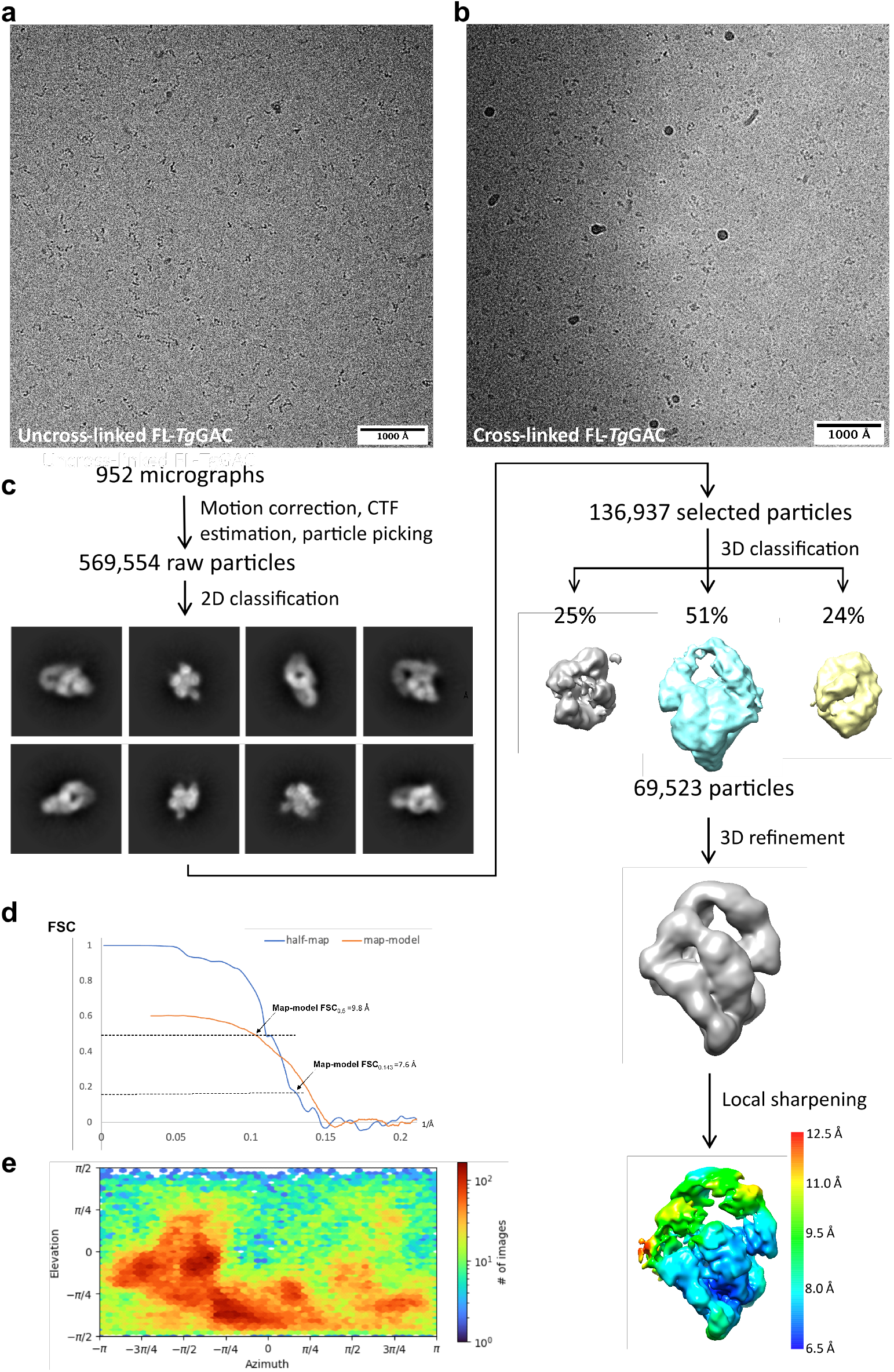
Workflow of single particle analysis of FL-*Tg*GAC. a. A typical field of view of native FL-*Tg*GAC. b. A typical field of view of crosslinked FL-*Tg*GAC. c. Flow chart for data processing of crosslinked FL-*Tg*GAC. Representative 2D classes are shown. The final refined map is based on 69523 particles. The local sharpened map is colour coded with local resolution. d. Gold standard Fourier shell correlation (FSC) and map-model FSC curves with estimated resolutions calculated in Phenix. Coil 1-2 of the crystal structure of FL-*Tg*GAC was used for map-model FSC. e. Angular distribution of the particles used in the final homogeneous refinement.

**Extended Data Fig. 4.**
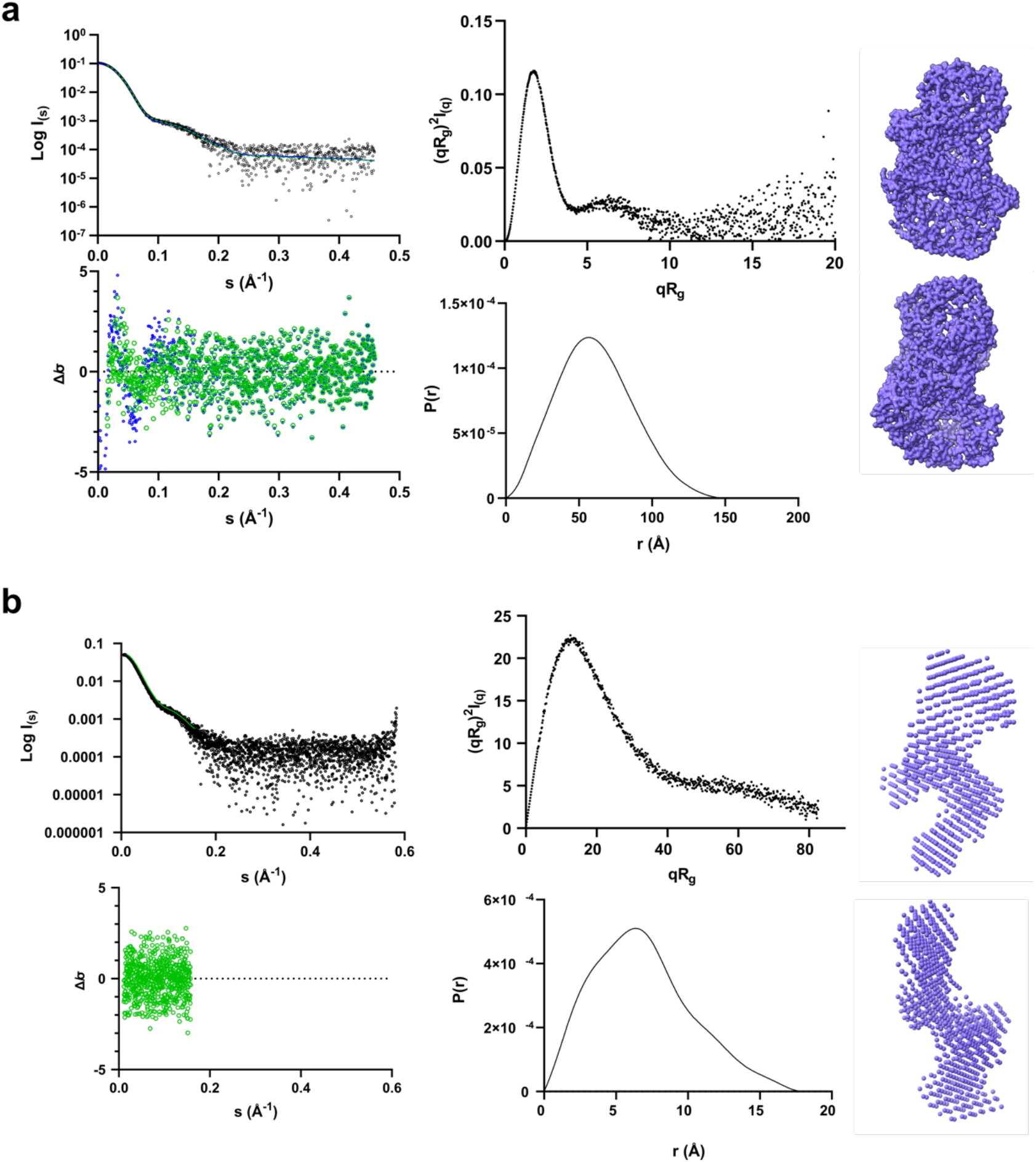
Conformations of FL-*Tg*GAC and *Tg*GAC coil 1-2-3 obtained from SEC-SAXS. a. Upper left: The fit of the GASBOR model of FL-*Tg*GAC (green line) and the crystal structure superimposed using CRYSOL (blue line) to the experimental SAXS data (black dots). The residuals are shown in the lower left panel, where Δ/σ= [I_exp_(q)-I_mod_(q)]/σ(q). The dimensionless Kratky plot (upper middle) suggests a folded globular protein. The distance distribution function (lower middle) suggests a globular protein with a maximum intramolecular distance of approximately 150 Å. On the right, an *ab initio* model generated using GASBOR is shown in two orientations. b. Upper left: The fit of the GASBOR model of *Tg*GAC coil 1-2-3 (green line) to the experimental SAXS data (black dots). The residuals are shown in the lower left panel, where Δ/σ= [I_exp_(q)-I_mod_(q)]/σ(q). The dimensionless Kratky plot (upper middle) suggests an elongated, partially unstructured flexible protein. The distance distribution function (lower middle) suggests the presence of several distinct domains with a maximum intramolecular distance of approximately 180 Å. On the right, an *ab initio* model generated using DAMMIN is shown in two orientations.

**Extended Data Fig. 5.**
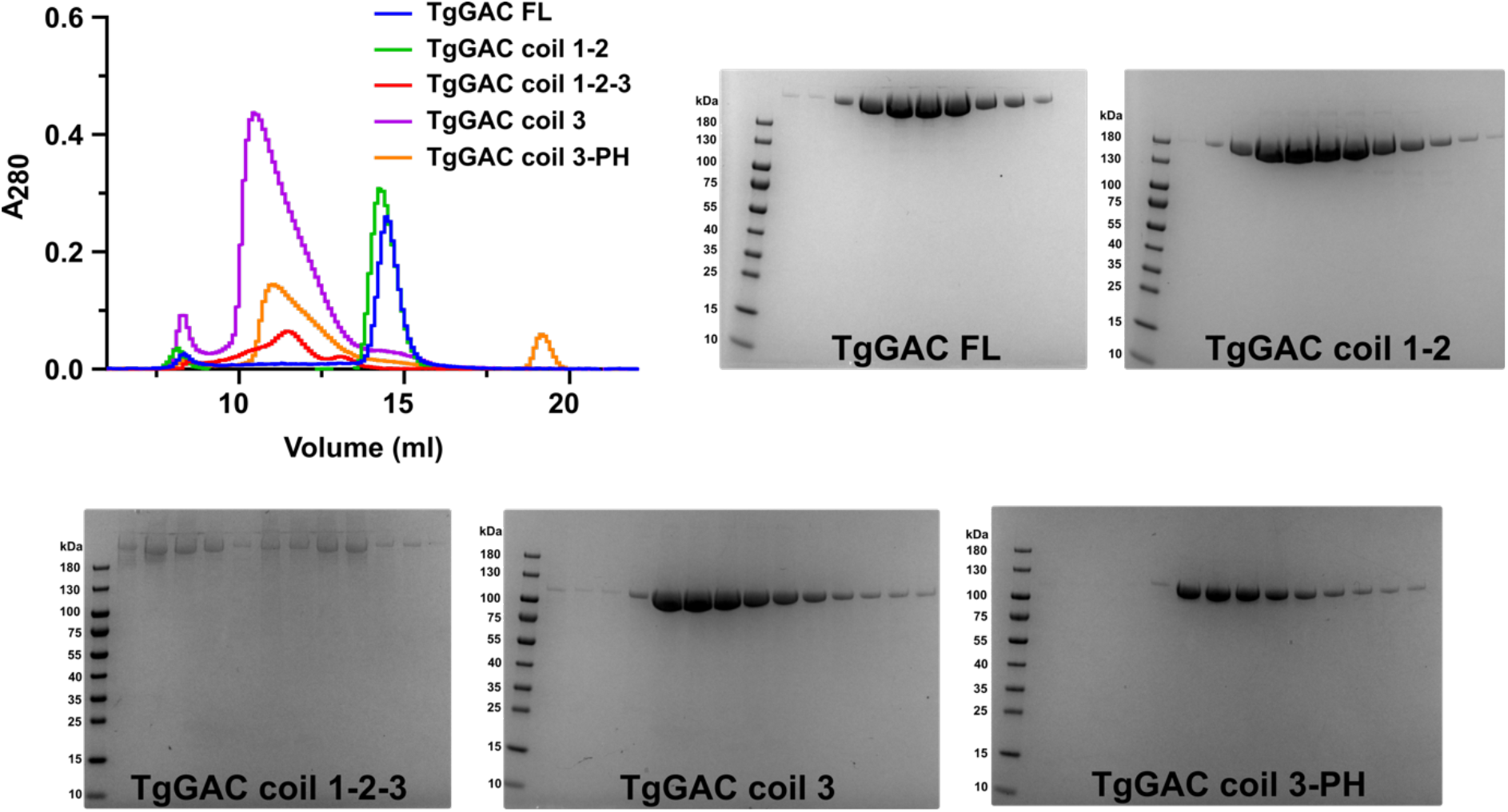
Analytical SEC of the *Tg*GAC subdomains. The SEC elution profiles of the different *Tg*GAC subdomains from a Superose 6 column is on the upper left. The curves correspond to FL-*Tg*GAC (black), coil 1-2 (green), coil 1-2-3 (red), coil 3 (blue), and coil 3-PH (purple). The respective Coomassie-stained SDS-PAGEs are shown in the other panels. The molecular weight marker sizes in kDa are indicated on the left side of each gel.

**Extended Data Fig. 6.**
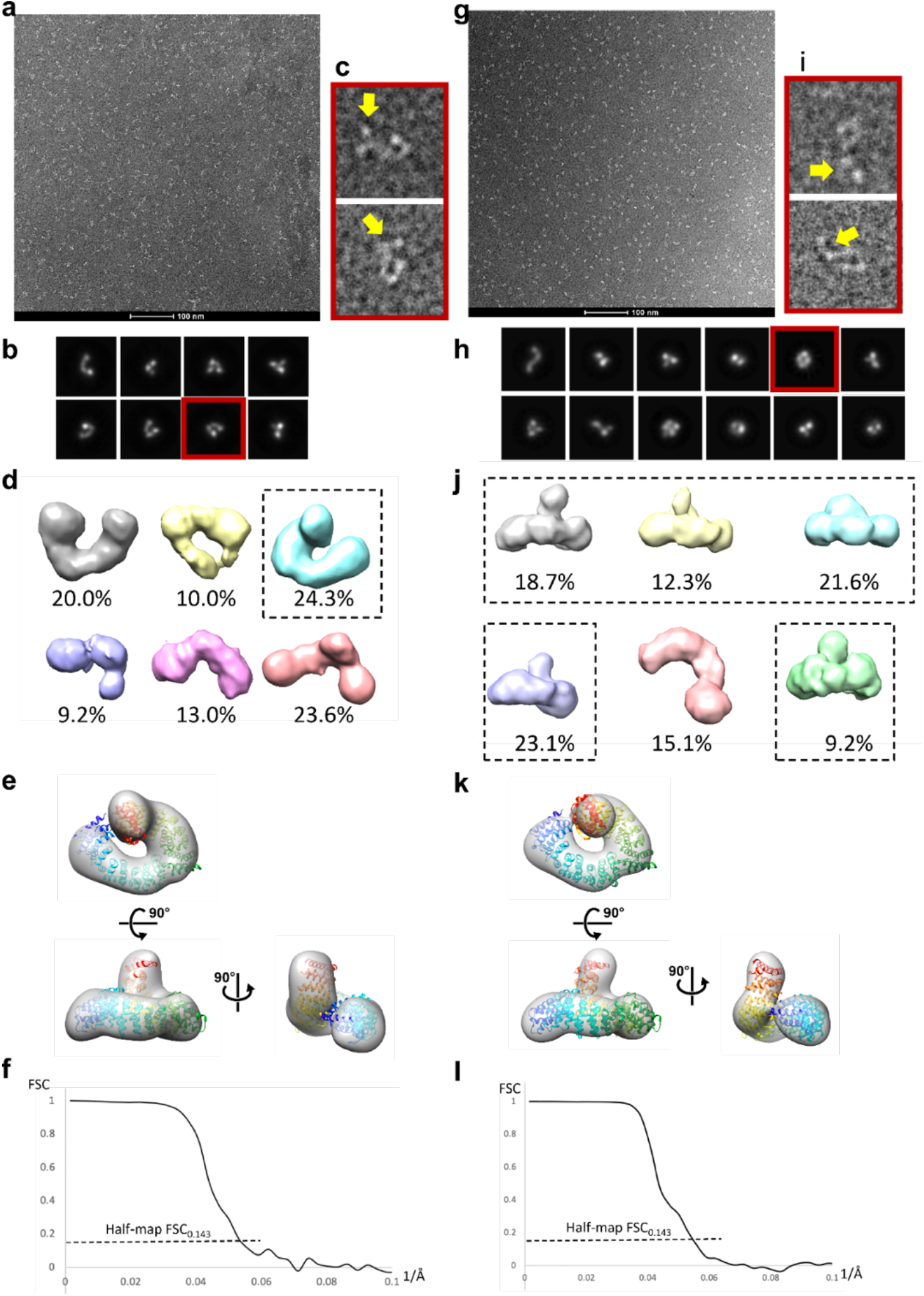
Negative stain imaging of *Tg*GAC coil 1-2 without (a-f) and with (g-l) 1% GA. a. A negative stain image of *Tg*GAC coil 1-2 (native). b. Representative 2D classes of native *Tg*GAC coil 1-2. c. Two images of individual particles within the 2D class highlighted in red in (b). The central regions of the images resemble the 2D class average, with extra densities likely to be coil 1 indicated by the yellow arrow. d. Results of 3D classification. The class in blue matches the structure of coil 2, which is then refined (e). f. Gold standard FSC of the map in (h). The resolution is approximately 20 Å. g. A negative stain image of *Tg*GAC coil 1-2 (crosslinked). h. Representative 2D classes of crosslinked *Tg*GAC coil 1-2. i. Two images of individual particles within the 2D class highlighted in red in (h). The central regions of the images resemble the 2D class average, with extra densities likely to be coil 1 indicated by the yellow arrow. j. Results of 3D classification. Five out of six classes (85%) match the structure of coil 2, which are then combined and refined into a single map (k). l. Gold standard FSC of the map in (k). The resolution is approximately 20 Å.

**Extended Data Fig. 7.**
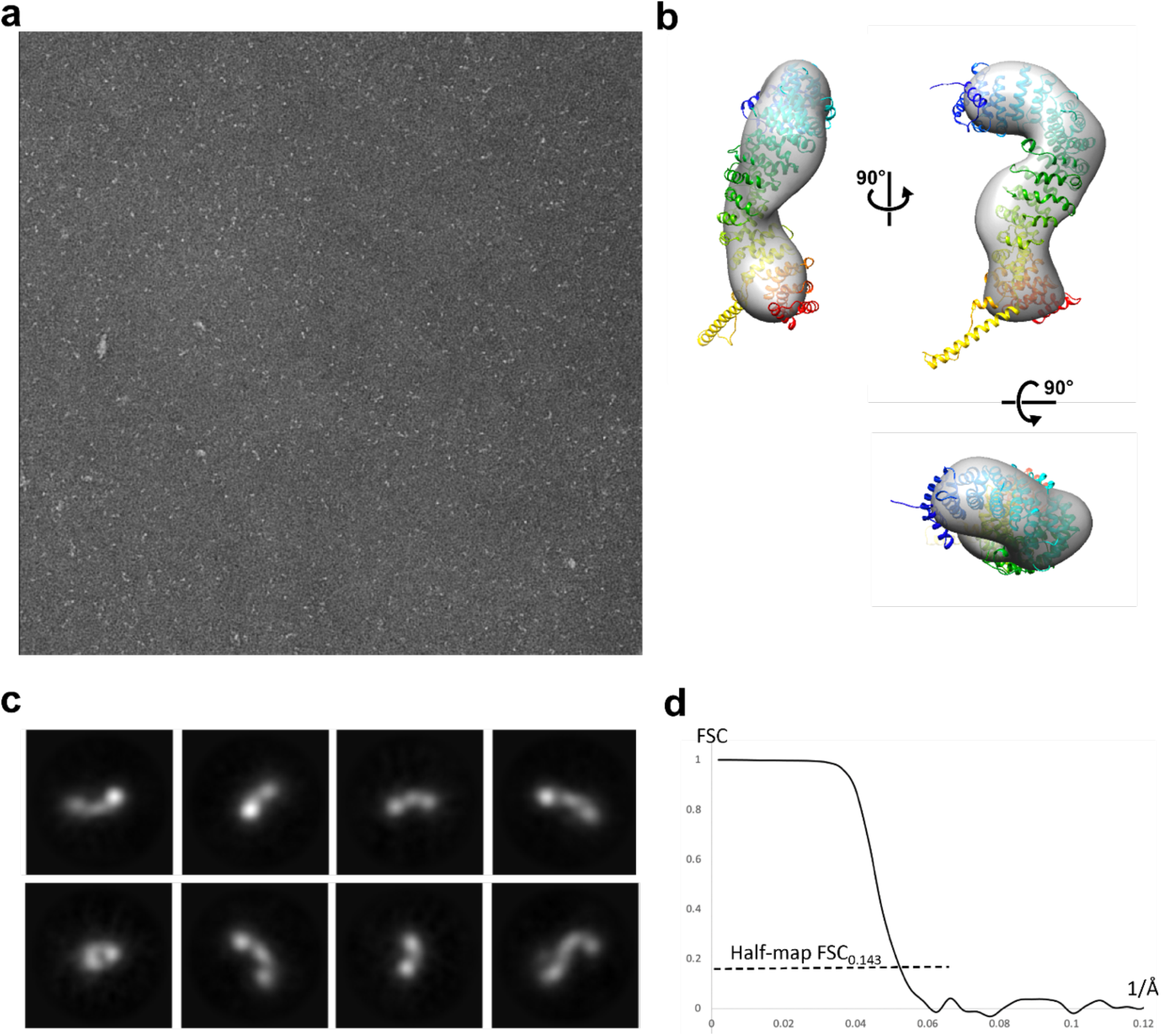
Negative stain structure of *Tg*GAC coil 3. a. A negative stain image of *Tg*GAC coil 3 (native). b. Representative averaged 2D classes showing different viewing angles. c. negative stain map of *Tg*GAC coil 3 with the crystal structure. (d) Gold standard FSC of the map in (c). The resolution is approximately 20 Å.

**Extended Data Fig. 8.**
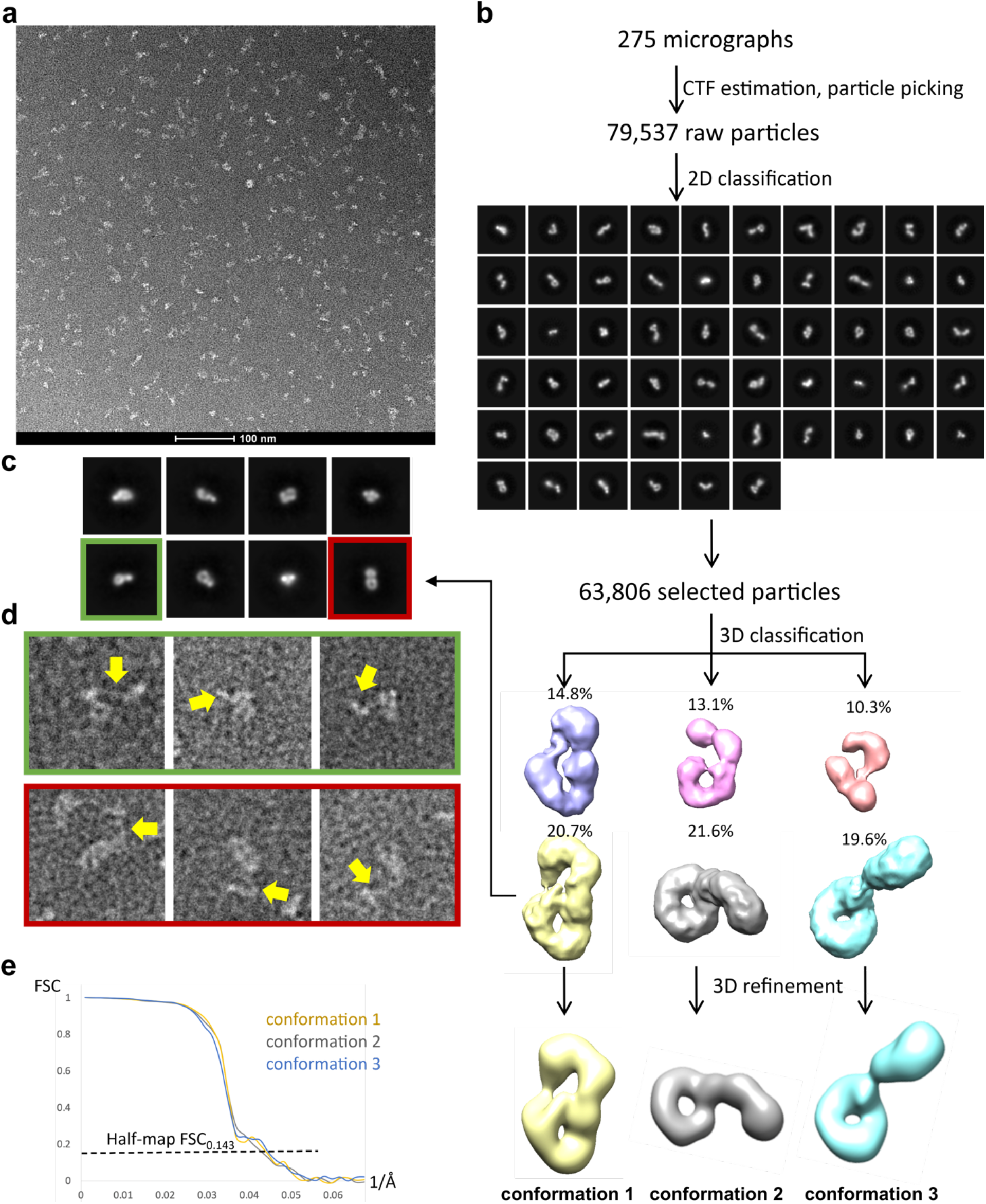
Workflow of single particle analysis of *Tg*GAC coil 1-3. a. A typical negative stain image of *Tg*GAC coil 1-3. b. Flow chart for data processing. Three classes from 3D classification were selected and refined respectively. c. A few 2D classes classified from the particles contributed to the 3D class in yellow. d. A few raw particle images from the two 2D classes highlighted in green and red. The central region of the images match the 2D class averages which represent the density of coil 2 and coil 3, while extra ‘tails’ extending from them are likely to be coil 1 (pointed by the yellow arrows). e. Gold standard Fourier shell correlation (FSC) of the three refined structures. The resolutions are all in the range of 20-25 Å.

**Extended data Fig. 9.**
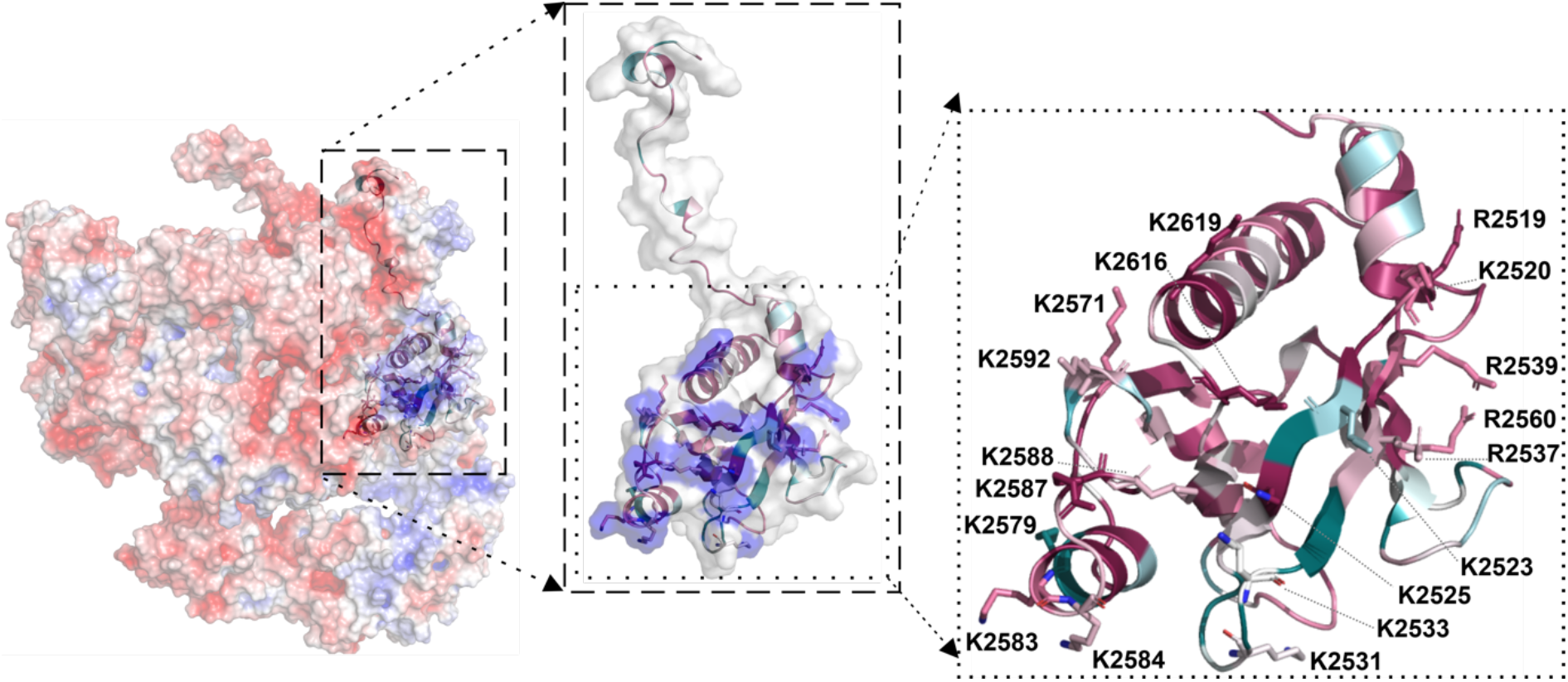
The *Tg*GAC PH domain has a solvent-accessible positively charged patch. The electrostatic surface potential of FL-*Tg*GAC is shown on the left. The insets show the zoomed-in surface and a cartoon representation of the PH domain with the 18 Lys residues highlighted in blue in the surface and as stick models and labeled in the cartoon representation.

## Notes

### Competing Interest Statement

The authors have declared no competing interest.

